# BRD2 inhibition blocks SARS-CoV-2 infection by reducing transcription of the host cell receptor ACE2

**DOI:** 10.1101/2021.01.19.427194

**Authors:** Avi J. Samelson, Quang Dinh Tran, Rémy Robinot, Lucia Carrau, Veronica V. Rezelj, Alice Mac Kain, Merissa Chen, Gokul N. Ramadoss, Xiaoyan Guo, Shion A. Lim, Irene Lui, James Nunez, Sarah J. Rockwood, Jianhui Wang, Na Liu, Jared Carlson-Stevermer, Jennifer Oki, Travis Maures, Kevin Holden, Jonathan S. Weissman, James A. Wells, Bruce R. Conklin, Benjamin R. TenOever, Lisa A. Chakrabarti, Marco Vignuzzi, Ruilin Tian, Martin Kampmann

## Abstract

SARS-CoV-2 infection of human cells is initiated by the binding of the viral Spike protein to its cell-surface receptor ACE2. We conducted a targeted CRISPRi screen to uncover druggable pathways controlling Spike protein binding to human cells. We found that the protein BRD2 is required for *ACE2* transcription in human lung epithelial cells and cardiomyocytes, and BRD2 inhibitors currently evaluated in clinical trials potently block endogenous *ACE2* expression and SARS-CoV-2 infection of human cells, including those of human nasal epithelia. Moreover, pharmacological BRD2 inhibition with the drug ABBV-744 inhibited SARS-CoV-2 replication in Syrian hamsters. We also found that BRD2 controls transcription of several other genes induced upon SARS-CoV-2 infection, including the interferon response, which in turn regulates the antiviral response. Together, our results pinpoint BRD2 as a potent and essential regulator of the host response to SARS-CoV-2 infection and highlight the potential of BRD2 as a novel therapeutic target for COVID-19.

## Introduction

The ongoing COVID-19 pandemic is a public health emergency. As of September 2021, SARS-CoV-2, the novel coronavirus causing this disease, has infected over 200 million people worldwide, causing at least four and a half million deaths (https://covid19.who.int). New infections are still rapidly increasing despite current vaccination programs. The emergence of novel viral variants with the potential to partially overcome vaccine-elicited immunity highlights the need to elucidate the molecular mechanisms that underlie SARS-CoV-2 interactions with host cells to enable the development of therapeutics to treat and prevent COVID-19, complementing ongoing vaccination efforts.

SARS-CoV-2 entry into human cells is initiated by the interaction of the viral Spike protein with its receptor on the cell surface, Angiotensin-converting enzyme 2 (ACE2). To uncover new therapeutic targets targeting this step of SARS-CoV-2 infection, we conducted a focused CRISPR interference (CRISPRi)-based screen for modifiers of Spike binding to human cells. We expected that ACE2 and factors regulating ACE2 expression would be major hit genes in this screen. A second motivation for identifying regulators of ACE2 was the fact that ACE2 affects inflammatory responses and is itself regulated in the context of inflammation^1–3^. Inflammatory signaling, in particular the type I interferon response, is known to be misregulated in the most severely affected COVID-19 patients^4–7^. Therefore, regulators of ACE2 expression would likely be relevant for COVID-19 in human patients, as suggested by clinical data^8^.

Previous CRISPR screens have been performed in cell-based models of SARS-CoV-2 infection that overexpressed an ACE2 transgene^9,10^, represented cell types not primarily targeted by SARS-CoV-2^11^, or were non-human cells^12^. While these studies elucidated major features of SARS-CoV-2 biology, we reasoned that the cell lines used would not have enabled the discovery of regulators of ACE2 expression in relevant human cell types.

Here, we selected a lung epithelial cancer cell line, Calu-3, which endogenously expresses ACE2, to perform a targeted CRISPRi screen to find novel regulators of Spike protein binding. We found that the strongest hit genes are potent regulators of ACE2 levels. Knockdown of these genes reduced or increased ACE2 levels transcriptionally, and prevented or enhanced, respectively, SARS-CoV-2 infection in cell culture.

We identified the transcriptional regulator Bromodomain-containing protein 2 (Brd2) as a major node for host-SARS-CoV-2 interaction. Brd2 is part of the Bromodomain and Extra-terminal domain (BET) family of proteins that includes Brd3, Brd4 and BrdT. BETs are being explored as targets for a number of cancers^13^. These proteins are known to be master transcriptional regulators and serve to bridge chromatin marks (mostly acetyl-lysines) to the transcriptional machinery^14^. We found Brd2 inhibition to downregulate ACE2 expression in Calu-3 cells, iPSC-derived cardiomyocytes, primary human lung epithelial cells and reconstructed human nasal epithelia. Inhibition of BRD2 with small molecules, some of which are in phase I clinical trials, inhibited SARS-CoV-2 infection in primary human nasal epithelia and in Syrian hamsters. We propose Brd2 as a key regulator and potential therapeutic target for COVID-19.

## Results

### CRISPRi Screen for determinants of Spike-RBD binding to human cells

To identify cellular mechanisms controlling the binding of SARS-CoV-2 to human cells, we identified a cell line that would robustly bind the viral spike protein (S). We measured binding of a previously described recombinant protein construct encompassing the SARS-CoV-2 Spike protein receptor-binding domain with a C-terminal human IgG Fc-domain fusion^15^, referred to hereafter as Spike-RBD, to several commonly used human cell lines (Fig. 1a and Extended Data Fig. 1a). Within this cell line panel, only Calu-3 cells displayed a binding curve consistent with specific binding of Spike-RBD, with an EC_50_ of 9.72 nM (95% CI: 4.37 – 22.42 nM). This value agrees with the dissociation constant of Spike-RBD-ACE2 binding determined *in vitro*, 4.7 nM^16^, within measurement error. Spike-RBD binding is dependent on ACE2 expression, as binding is abrogated in Calu-3 *ACE2* knockout cells (Fig. 1b). Calu-3 cells are challenging to culture compared to most cell lines - they proliferate slowly, and their adherence properties pose challenges for flow cytometry. Nevertheless, Calu-3 cells are particularly attractive cell culture model for studying SARS-CoV-2 from the biological point of view, because they are derived from lung epithelia, which is selectively infected by SARS-CoV-2 in patients^17^, are known to support infection of SARS-CoV and SARS-CoV-2^18^ and have recently been reported to closely recapitulate gene expression changes that occur in patients^19^.

**Figure 1:**
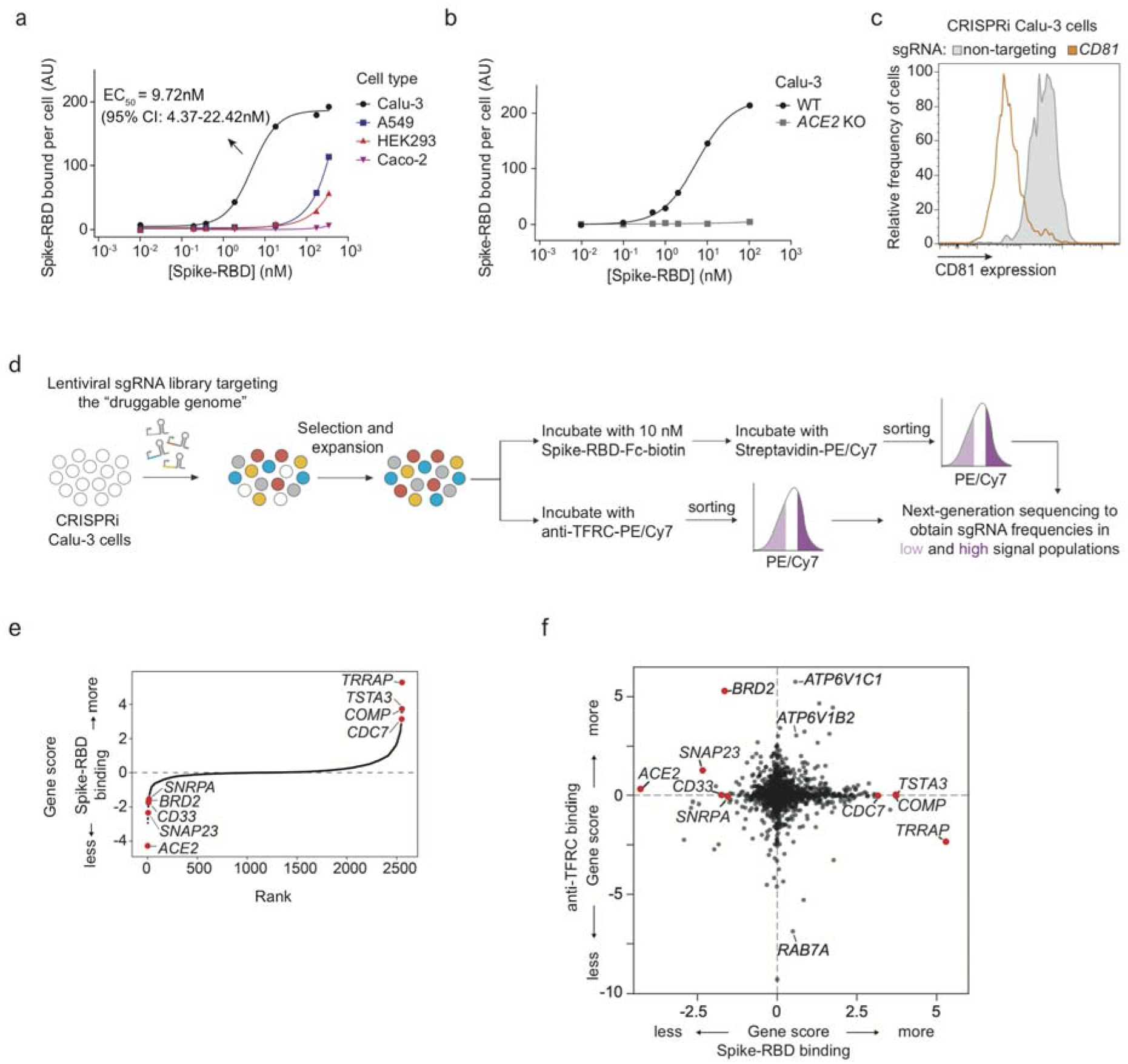
CRISPRi screen reveals cellular factors controlling Spike protein binding. **a**, Human cell lines were incubated with different concentrations of recombinant SARS-CoV-2 Spike protein receptor-binding domain (Spike-RBD) and the amount of Spike-RBD bound per cell was quantified by flow cytometry. Only Calu3 cells (black) display saturable binding, for which an EC_50_ value was fit. Error of EC_50_ is the 95% confidence interval. **b**, Spike-RBD binding is eliminated in Calu-3 cells with ACE2 knockout (grey). **c**, Validation of CRISPRi activity of a Calu-3 CRISPRi line. Calu-3 cells stably expressing CRISPRi machinery were transduced with an sgRNA targeting CD81 (orange) or a non-targeting sgRNA (grey), and CD81 levels were determined by flow cytometry. **d**, CRISPRi screen strategy. CRISPRi Calu-3 cells transduced with an sgRNA library targeting the “druggable genome” were stained either with Spike-RBD or an anti-TFRC antibody. Cells were then FACS-sorted into bins (top and bottom 30%) based on Spike-RBD or anti-TFRC binding signal, and frequencies of cells expressing each sgRNA were determined for each bin by targeted next-generation sequencing. **e**, Rank-order plot of Spike-RBD hit genes. Genes selected for follow-up experiments are highlighted as red dots. **f**, Scatter plot of gene scores for the Spike-RBD screen (x-axis) and the anti-TFRC screen (y-axis). Genes selected for follow-up experiments are highlighted as red dots.

We generated a polyclonal Calu-3 line constitutively expressing machinery to enable CRISPRi-based genetic screens^20,21^ and validated its CRISPRi activity (Fig. 1c and Extended Data Fig. 1b). CRISPRi uses a catalytically dead Cas9 (dCas9) fused to a transcriptional repressor, KRAB, to knockdown genes at specific sites programmed by the loading of the dCas9-KRAB with a single guide RNA (sgRNA). Using this line, we then performed a focused CRISPRi screen for factors controlling Spike-RBD binding (Fig. 1d). In order to maximize our chances of identifying potential novel therapeutic targets for COVID-19, we screened a sgRNA library targeting the “druggable genome”, comprising ~2,300 genes, with ~16,000 total sgRNAs including non-targeting control sgRNAs^22^. In parallel, we screened the same library using a fluorophore-conjugated antibody against the transferrin receptor (TFRC, also known as CD71), to control for factors that generally affect protein trafficking or protein binding to the cell surface (Fig. 1d–f). Due to the limitations of the Spike-RBD binding assay and Calu-3 cells, this screen was conducted at a lower average representation (~100 sorted cells / sgRNA) than ideal, resulting in relatively high noise and therefore fewer hits crossing a false discovery rate cutoff of 0.1 than in typical CRISPR screens (see Extended Data Figure 2 and Methods).

Despite the increased noise, *ACE2*, as expected, was the strongest hit gene, knockdown of which decreased binding of Spike-RBD, while having no effect on TFRC levels (Fig. 1e,f). Conversely, *RAB7A*, which was recently reported to be essential for the trafficking of TFRC to the cell surface^23^, was the strongest hit that decreased TFRC levels, with no effect on Spike-RBD binding (Fig. 1f). Generally, hits were not correlated between the two screens (Fig. 1f), demonstrating the specificity of each screen. While the screens did not result in a large number of strong hits (Extended Data Table 1), we decided to validate the top 15 genes knockdown of which decreased Spike-RBD binding and the top five genes knockdown of which increased Spike-RBD binding. We cloned individual sgRNAs targeting each of these genes and evaluated their effect on Spike-RBD binding (Extended Data Figure 3). Based on these experiments, we selected hits that robustly recapitulated their phenotypes from the primary screen for further characterization: two genes knockdown of which decreased Spike-RBD binding (*ACE2* and *BRD2*), and three genes knockdown of which increased Spike-RBD-binding (*CDC7*, *COMP* and *TRRAP*).

### Hit genes modulate ACE2 levels and affect infection with SARS-CoV-2

Since Spike-RBD binding is dependent on ACE2 expression as shown above, we hypothesized that other hit genes might act by modulating ACE2 levels. Western Blots for ACE2 levels in Calu-3 cell lines expressing sgRNAs against validated target genes (hereafter referred to as knockdown lines) indeed revealed marked changes in ACE2 protein levels. For hits associated with lower levels of Spike-RBD binding in the primary screen, we observed lower levels of ACE2 protein, and vice-versa for those hits associated with higher levels of Spike-RBD binding (Fig. 2a,b). To distinguish whether hit genes affected ACE2 protein levels via transcriptional or post-transcriptional mechanisms, we performed qPCR to measure *ACE2* transcript levels in these same knockdown lines. For all tested genes, we observed changes in *ACE2* transcript levels that were concordant with the changes in ACE2 protein levels (Fig. 2c), indicating that they acted on the transcriptional level. Some genes, such as *COMP* and *TRAPP*, showed relatively modest effects on *ACE2* transcript levels, but quite large effects on ACE2 protein levels, suggesting that these hit genes additionally affect post-transcriptional regulation of ACE2 expression.

**Figure 2:**
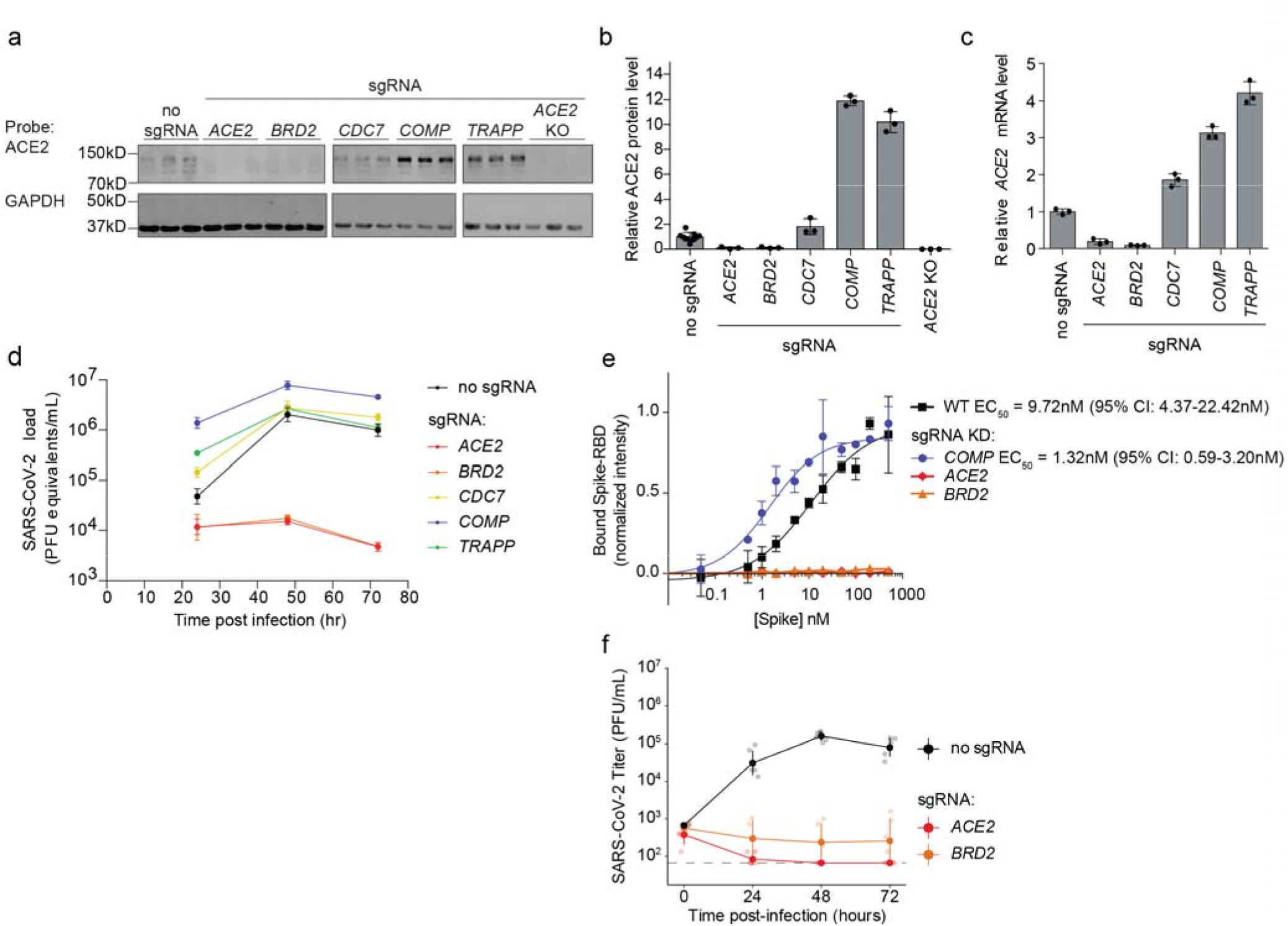
Hit genes modulate ACE2 levels and SARS-CoV-2 infection. **a**, Western blotting for ACE2 and GAPDH in Calu-3 CRISPRi cells expressing no sgRNA or sgRNAs targeting different hit genes, or ACE2 knockout Calu-3 cells. Three lanes represent biological triplicates for each cell line. **b**, Quantification of ACE2 protein levels relative to GAPDH based on the data in (a). Average and standard deviation for three biological replicates are shown. **c**, Relative amounts of ACE2 transcript levels measured by qPCR in Calu-3 CRISPRi cells expressing sgRNAs targeting different hit genes, compared to cells without sgRNA. Average and standard deviation for three technical replicates are shown. **d**, Calu-3 CRISPRi cells expressing different sgRNAs targeting hit genes were infected with SARS-CoV-2 and viral RNA in the supernatant measured by RT-qPCR as a function of time post-infection. Average and standard deviation of three wells are shown. **e**, Spike-RBD binding to Calu-3 cells was quantified by flow cytometry of Calu3 cells expressing sgRNAs targeting individual hit genes after incubation with increasing concentrations of Spike-RBD. For genes for which data could be fitted with a binding curve, the EC_50_ was determined along with the 95% confidence intervals. Data points are average values from three biological replicates for each gene knockdown with error bars representing the standard deviation, except for *ACE2* and *BRD2* where only one experiment at each concentration was performed. **f**, Plaque assays in Calu-3 CRISPRi cells expressing different sgRNAs targeting hit genes were infected with SARS-CoV-2 as a function of ti e post-infection. Average and standard deviation of six biological replicates are shown.

We next determined the effect of hit gene knockdown on susceptibility to SARS-CoV-2 infection. We infected cells expressing sgRNAs against hit genes with SARS-CoV-2 and measured virus replication 24, 48 and 72 hours post-infection using RT-qPCR (Fig. 2d). Already at 24 hours post-infection, viral genome copies diverged concordantly with changes in ACE2 levels and Spike-RBD binding: sgRNAs that lowered Spike-RBD binding reduced virus replication, while sgRNAs that increased Spike-RBD binding resulted in higher virus replication. *BRD2* knockdown abrogated viral replication in these cells to similar levels as ACE2 knockdown, even at 72 hours post-infection, while *COMP* knockdown supported an order of magnitude increase in viral titers. Focusing on these three hit genes, *ACE2*, *BRD2* and *COMP*, we then quantified how gene knockdown modulates Spike-RBD binding to cells (Fig 2e). Knockdown of *ACE2* and *BRD2* abolished spike binding, while *COMP* decreased the EC50 compared to WT from 9.72 nM (95% CI: 4.37 – 22.42 nM) to 1.32nM (95% CI: 0.59 – 3.20 nM), by almost a full order of magnitude (Fig. 2e). We also confirmed that gene knockdown decreased SARS-CoV-2 replication by performing plaque assays on WT, *BRD2* KD, and *ACE2* KD cells (Fig 2f).

### BRD2 inhibitors prevent SARS-CoV-2 infection of human cells

Given the stringent inhibition of SARS-CoV-2 infection achieved by BRD2 knockdown, and the fact that BRD2 is currently being evaluated as therapeutic target in cancer^13,24^, with several small molecule inhibitors in clinical trials^25^, we decided to focus on this hit gene.

We validated that CRISPRi knockdown of *BRD2* robustly reduced Brd2 protein levels (Extended Data Fig. 4). Transgenic expression of full-length Brd2 restored *ACE2* transcript levels (Fig. 3a), validating that the reduction in ACE2 expression triggered by CRISPRi targeting of BRD2 was indeed due to BRD2 knockdown. Transgenic expression of truncation mutants of BRD2 did not rescue ACE2 expression (Fig. 3a), indicating that full-length Brd2 is required for ACE2 expression.

**Figure 3:**
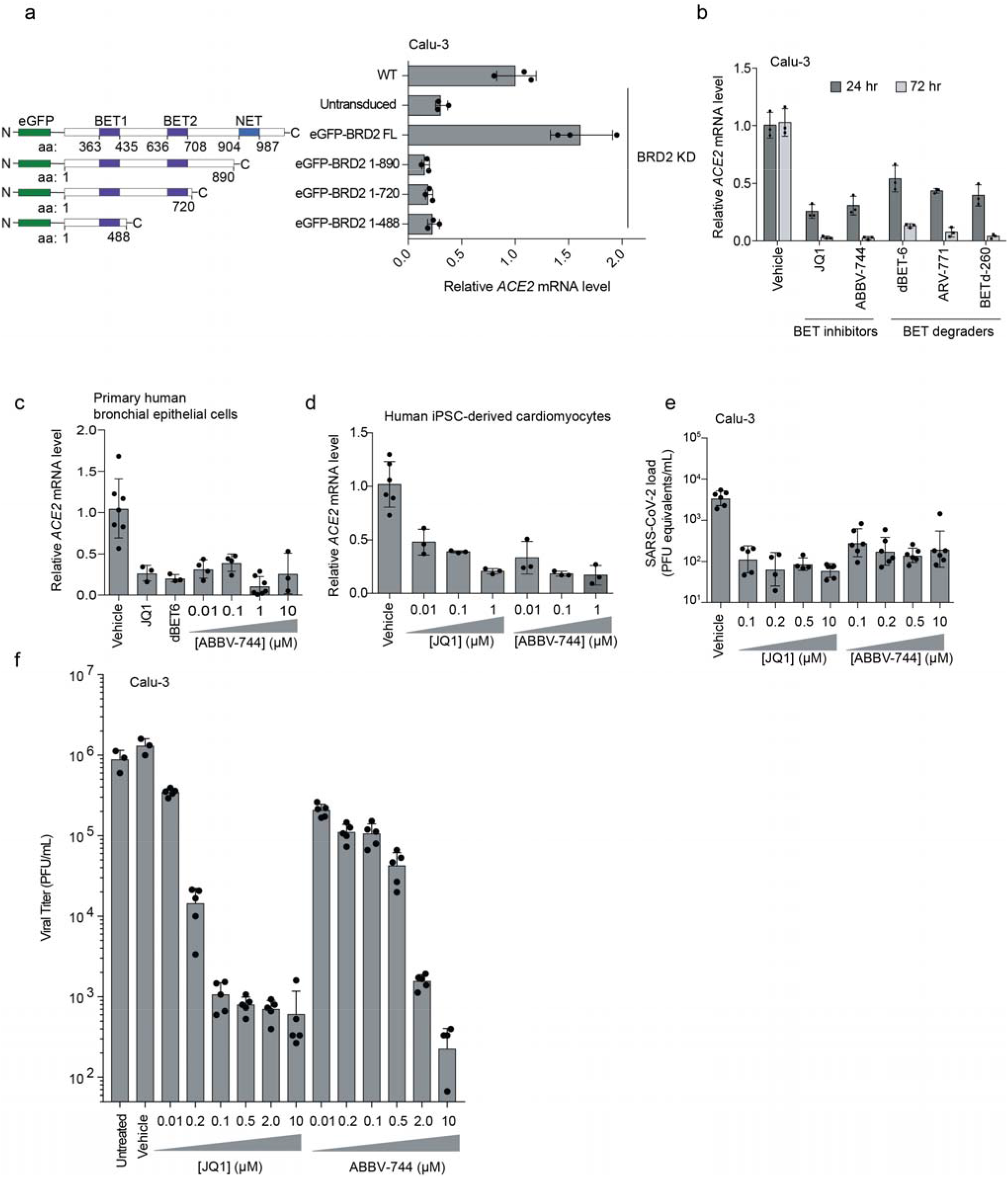
BRD2 inhibitors potently reduce ACE2 levels and SARS-CoV-2 infection. **a**, Transgenic constructs expressed in Calu-3 cells (left). Transcript levels of ACE2 relative to ACTB in Calu3 cells transduced with eGFP-BRD2 truncations in a BRD2 knockdown background (right). ACE2 expression is relative to WT. Average and standard deviation of technical triplicates are shown for each transduced construct. **b**, Transcript levels of ACE2 relative to ACTB in Calu3 cells treated with BRD2 inhibitors (JQ1 at 10 μM, ABBV-744 at 10 μM and dBET-6 at 200 nM) were quantified at 24 (dark grey bars) and 72 hours (light grey bars) post-treatment. Average and standard deviation of technical triplicates are shown for each condition. **c**, Transcript levels of ACE2 relative to ACTB in primary human bronchial epithelial cells treated (NHBE) with BRD2 inhibitors (JQ1 at 10 μM, dBET-6 at 20 nM, ABBV744 at 0.01-10uM) were quantified at 72 hours post-treatment. Average and standard deviation of biological triplicates are shown for each condition. **d**, Transcript levels of ACE2 relative to 18S rRNA in human iPSC-derived cardiomyocytes treated with the indicated concentrations of BRD2 inhibitors were quantified at 72 hours post-treatment. Average and standard deviation of 3 or more biological replicates are shown for each condition. **e**, SARS-CoV-2 viral RNA in the supernatant measured by RT-qPCR 24 hours post-infection of Calu-3 cells infected 72 hours after treatment with the indicated concentrations of BRD2 inhibitors. Average and standard deviation of four or more biological replicates are shown for each condition. **f,** Plaque assays in Calu-3 CRISPRi cells treated with increasing concentrations of the BET inhibitors JQ-1 or ABBV-744 infected with SARS-CoV-2 as a function of time post-infection. Average and standard deviation of six biological replicates are shown except for vehicle and untreated which are three biological replicates.

To test the potential of Brd2 as a therapeutic target for COVID-19, we treated cells with a panel of compounds targeting BRD2: two BET domain inhibitors, (JQ1^26^ and ABBV-744^27^, which is currently in clinical trials NCT03360006 and NCT04454658), and three Proteolysis Targeting Chimeric (PROTAC) compounds that lead to the degradation of Brd2 (dBET-6^28^, ARV-771^29^, and BETd-260^29^). After only 24 hours of treatment with these drugs, *ACE2* mRNA levels measured by qPCR decreased roughly two-fold (Fig. 3b). This effect was magnified after treatment for 72 hours, when almost no *ACE2* mRNA was detectable for any of the Brd2-targeting compounds tested, phenocopying Brd2 knockdown (Fig. 3b). Similarly, we found that BET inhibitors led to substantial decreases in *ACE2* mRNA levels in primary human bronchial epithelial cells (Fig. 3c) and human iPSC-derived cardiomyocytes (Fig. 3d), two non-transformed cell types that are susceptible to SARS-CoV-2 infection^30,31^. Importantly, BET inhibitors were non-toxic to Calu-3 cell, primary human bronchial epithelial cells, and cardiomyocytes at effective concentrations (Extended Data Fig. 5).

Since pharmacological inhibition of Brd2 phenocopied *BRD2* knockdown, we hypothesized that these same compounds might prevent infection of cells exposed to SARS-CoV-2. To test this, we treated Calu-3 cells for 72 hours with the BET inhibitors JQ-1 and ABBV-744, and measured SARS-CoV-2 replication at 48 hours post infection. Strikingly, we found that treated cells displayed 100-fold decreased viral replication versus untreated cells (Fig. 3e), a similar effect size compared to *BRD2* or *ACE2* knockdown (Fig. 2d,e).

### BRD2 regulates the transcription of ACE2 and other host genes induced by SARS-CoV-2 infection

We next asked whether Brd2 controls transcription of additional genes beyond *ACE2*. We performed RNA sequencing of Calu-3 cells after treatment with the BET-domain inhibitors JQ-1 and ABBV-744 as well as *BRD2* CRISPRi knockdown (Extended Data Table 2). We also included CRISPRi knockdown of two other validated hit genes from our screen, *COMP* and *ACE2* as well as over-expression of the viral protein E, which had been reported to interact with Brd2^32^. RNA-seq of BRD2 knockdown and BET domain inhibitor treated cells recapitulated downregulation of *ACE2* (Fig. 4a). *TMPRSS2,* the gene encoding a protease important for viral entry in many cell types, was not a differentially expressed gene in any condition (Extended Data Table 2). Surprisingly, *BRD2* knockdown or pharmacological inhibition also resulted in marked downregulation of genes involved in the type I interferon response, while *ACE2* knockdown slightly increased expression of those same genes (Fig. 4a,b). Furthermore, the genes downregulated by both BRD2 knockdown and inhibition were strongly enriched in genes induced by SARS-CoV-2 infection in patient and cultured cells (Fig. 4c).

**Figure 4:**
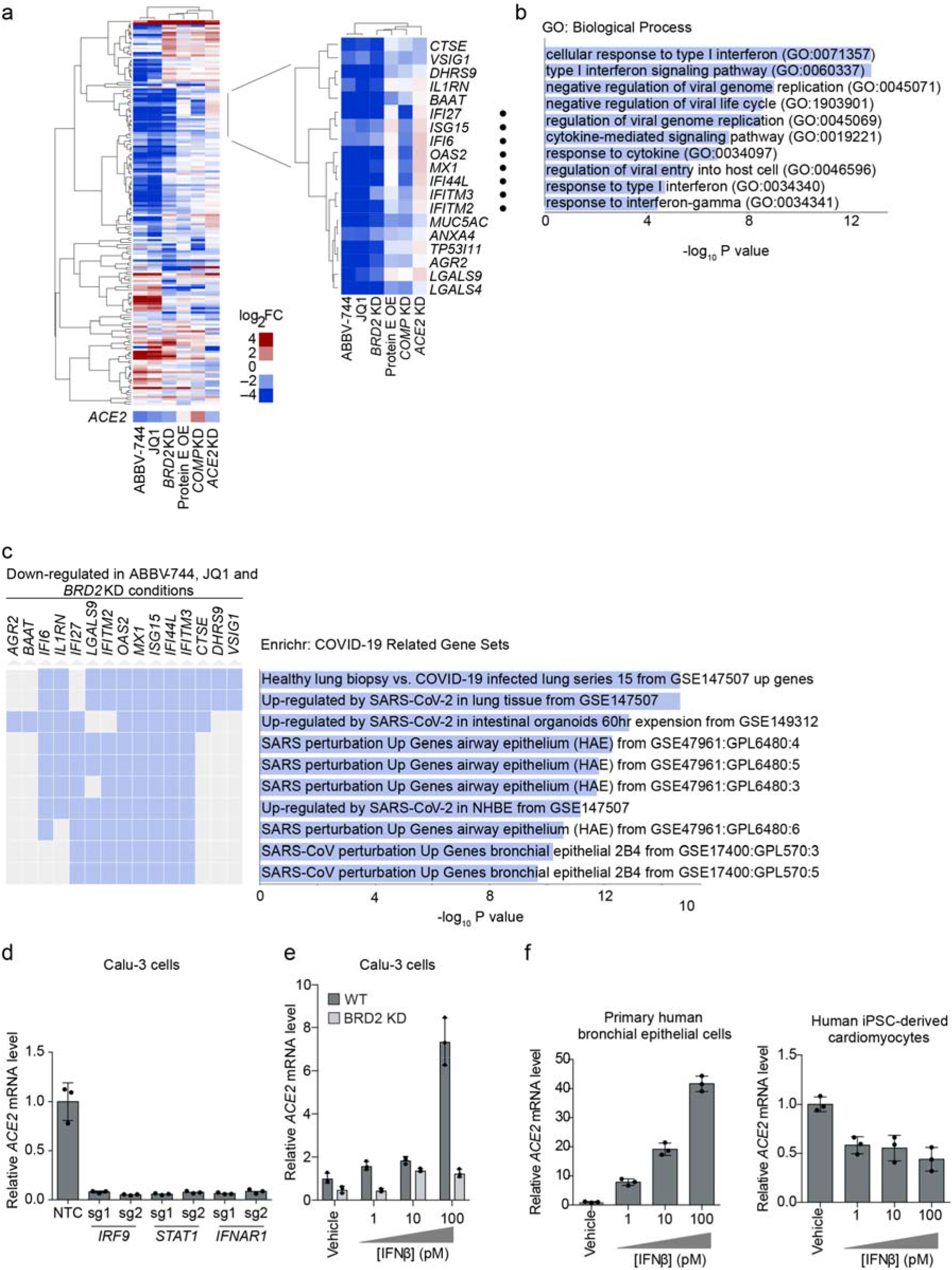
BRD2 controls genes induced by interferon and SARS-CoV-2 infection. **a**, Differentially expressed genes from RNA sequencing of Calu-3 cells under different treatment conditions compared to control cells: 72-hour treatment with 10 μM JQ1 or 10 μM ABBV-744, *BRD2* knockdown (KD), SARS-CoV-2 protein E overexpression (OE), *COMP* KO, *ACE2* KD. Experiments were conducted in the absence of virus or interferon treatment. Heatmap showing log_2_-fold change (log_2_FC) for each condition relative to untreated controls (columns) for genes that are among top 50 differentially expressed genes (ranked by P values) in at least one of the conditions (rows). *ACE2* was not among these genes and is shown as a separate row. *Insert*, a cluster of genes that are down-regulated in both BRD2 inhibition by JQ1 and ABBV-744 and *BRD2* knockdown. Among these, genes associated with the GO term “Cellular Response to Type I interferon” are marked by black dots. **b**, Significantly enriched (FDR < 0.05) GO biological process terms for the genes shown in the inset in (a). **c**, Enrichment analysis for genes in the inset in (a) reveals COVID-19 related gene sets. Genes that appear in a gene set are marked in blue. **d**, Calu-3 cells expressing sgRNAs knocking down genes essential for interferon signal transduction assayed for transcript levels of *ACE2* relative to *ACTB* by qPCR. Average and standard deviation of 3 biological replicates are shown for each condition. **e**, WT (dark grey) or BRD2 knockdown (light grey) Calu-3 cells were treated with the indicated concentrations of interferon-beta (IFNβ), and transcript levels of *ACE2* relative to *ACTB* were quantified at 72 hours post-treatment by qPCR. Average and standard deviation of 3 biological replicates are shown for each condition. **f,** Primary human bronchial epithelial cells (left) and human iPSC-derived cardiomyocytes (right) were treated with the indicated concentrations of interferon-beta (IFNβ), and transcript levels of ACE2 relative to ACTB (for Calu-3 and primary human bronchial epithelial cells) or 18S rRNA (cardiomyocytes) were quantified at 72 hours post-treatment by qPCR. Average and standard deviation of 3 biological replicates are shown for each condition.

These findings are compatible with two distinct mechanisms: Brd2 could independently regulate *ACE2* and SARS-CoV-2-induced interferon response genes, or Brd2 could mediate the response to interferon, which in turn regulates *ACE2* transcription. *ACE2* expression has been reported to be induced by interferons in some studies^1,2^. Other studies, however, suggest that interferon suppresses *ACE2* expression^3^.

In Calu-3 cells, disruption of basal interferon signaling, via knockdown of the genes essential for interferon signal transduction *IRF9*, *STAT1*, or *IFNAR1*, abrogated *ACE2* expression (Fig. 4d, Extended Data Fig. 6). Conversely, treatment with exogenous IFNβ stimulates *ACE2* expression in a concentration dependent manner. Upon *BRD2* knockdown, however, this concentration-dependent increase in *ACE2* expression is inhibited (Fig. 4e). Thus, BRD2 is required for interferon-induced *ACE2* expression. Treatment with IFNβ similarly strongly increased *ACE2* mRNA levels in primary human bronchial epithelial cells, but reduced *ACE2* mRNA levels in human iPSC-derived cardiomyocytes (Fig. 4f), suggesting that the effect of interferons on *ACE2* can be context-dependent.

To test if Brd2 is a direct transcriptional regulator of *ACE2*, we performed CUT&RUN^33^ to comprehensively map genomic loci bound by Brd2 in Calu-3 cells (Extended Data Table 3). CUT&RUN is similar to ChIP-seq, as it measures the occupancy of factors bound to DNA, but has the advantage of higher sensitivity and lower requirement for cell numbers^33^. Genes adjacent to Brd2-bound sites detected in our experiment showed a highly significant overlap with Brd2-bound sites previously mapped by ChIP-seq in NCI-H23^34^ cells, another lung epithelium-derived cancer cell line (Fig. 5a). To further validate our CUT&RUN analysis, we performed Binding and Expression Target Analysis^35^ (BETA) to uncover direct BRD2 targets that were differentially expressed upon BRD2 knockdown, and identified several interferon response genes as direct BRD2 targets that were down-regulated (Fig. 5b). We verified that a previously described^34^ Brd2 binding side upstream of *PVT1* was also detected in our experiment (Fig. 5c). We also mapped a Brd2 binding site upstream of the several interferon-stimulated genes (ISGs), including *IRF9*, *STAT1*, and *MX1* (Fig. 5d). While there was some signal in the WT background at the *ACE2* locus that is decreased in *BRD2* knockdown cells, there were no peaks as determined by the peak calling algorithm (Fig 5e), suggesting that Brd2 is not a direct transcriptional regulator of *ACE2* expression.

**Figure 5:**
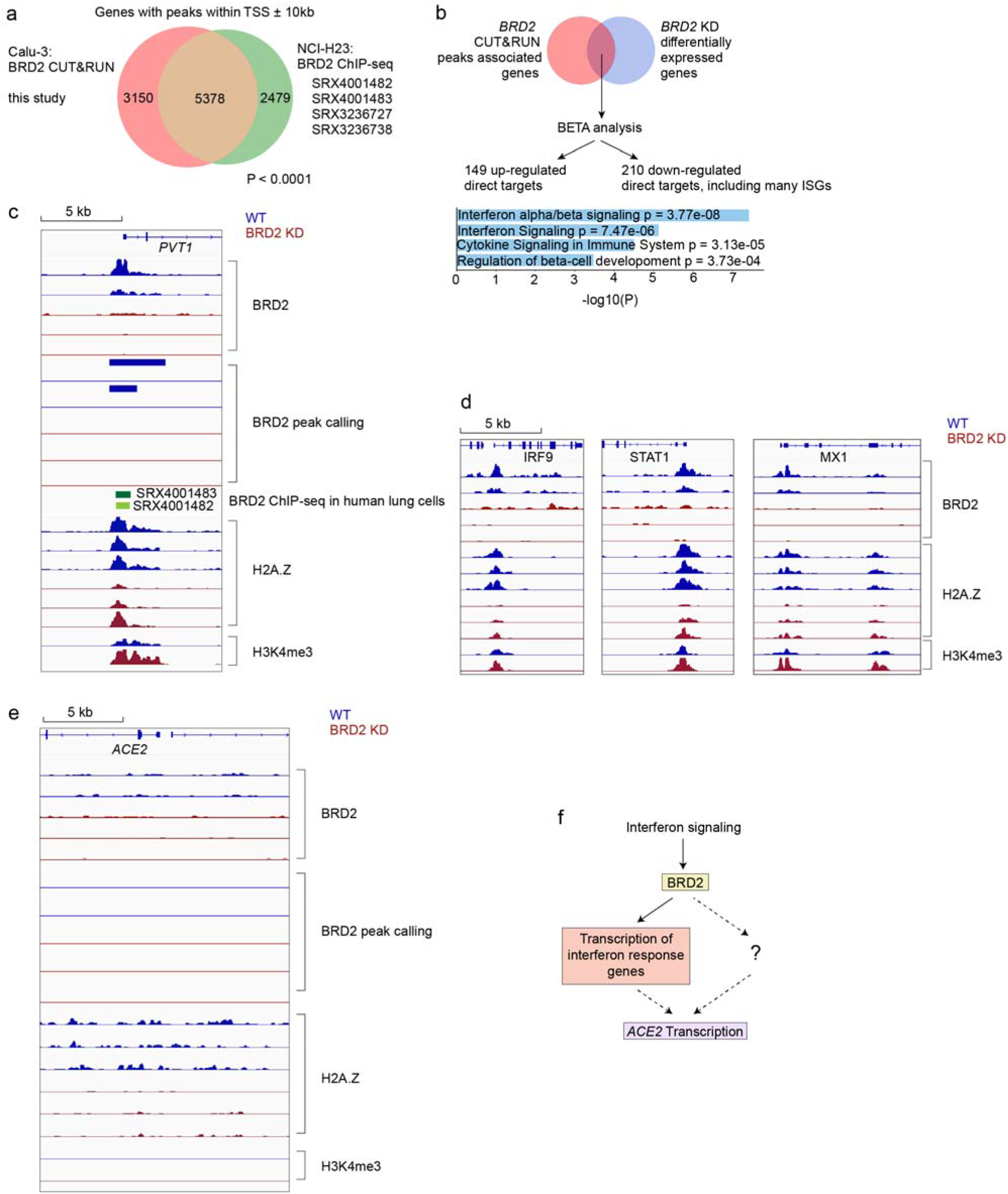
BRD2 directly regulates transcription of interferon-induced genes. **a**, Genes associated with BRD2 CUT&RUN peaks within 10kb of a transcription start site determined in this study in Calu-3 cells overlap significantly with published BRD2 ChIP-seq peaks from the indicated datasets (P < 0.0001, Fisher’s exact test). **b**, Binding and Expression Target Analysis (BETA) was performed to identify direct BRD2 targets that were differentially expressed upon *BRD2* knockdown. Many interferon response genes were identified as direct BRD2 targets. Direct BRD2 targets that were downregulated upon BRD2 knockdown were analyzed by ENRICHR for enriched Reactome pathways. Pathways with adjusted p-values less than 0.05 are displayed. **c-e** CUT&R experiments were conducted to map BRD2, H2A.Z and H3K4me3 genomic localization in WT (blue traces) and BRD2 KD (red) Calu-3 cells. Each trace represents an independent biological replicate. **c**, Known BRD2 regulatory sites are recapitulated. Raw signal tracks for WT and *BRD2* knockdown cells are shown at the known *BRD2* locus PVT1. *BRD2* CHIP-seq tracks from human lung cells are shown. *BRD2* peaks were called over IgG using SEACR at FDR < 0.05. **d**, Identified BRD2 and Histone H2A.Z occupancy and peak calling at ISGs in *BRD2* KD and WT Calu-3 cells. Raw signal tracks for BRD2, Histone H2A.Z, and H3K4me are shown. BRD2 peaks were called over IgG using SEACR at FDR < 0.05. BRD2 CHIP-seq tracks from human lung cells are also shown. **e**, Raw BRD2 signal tracks and peak calling as for c, at the *ACE2* locus. **f**, Proposed model for BRD2 control of ACE2 expression.

We also performed CUT&RUN for histone H2A.Z, which was previously reported to modulate the magnitude of ISG expression and thus connect Brd2 activity to interferon stimulation^36^. We found decreased H2A.Z occupancy at ISGs in *BRD2* knockdown cells (Fig. 5d), recapitulating the role of Brd2 as a potential chaperone of H2A.Z. These results support a model in which BRD2 controls the transcription of key interferon response genes, which can in turn induce *ACE2* transcription in some cell types (Fig 5f). Alternatively, *ACE2* expression may controlled by other genes that are expressed in a Brd2-dependent, interferon-stimulated manner (Fig. 5f).

### Brd2 inhibitors rescue cytotoxicity and reduce SARS-CoV-2 infection in human nasal epithelia and inhibit SARS-CoV-2 infection in Syrian hamsters

We then tested if ABBV-744, a Bromodomain inhibitor in clinical trials, could reduce SARS-CoV-2 infection and infection-associated phenotypes in more physiological models.

First, we investigated a human nasal epithelial model^37^. We treated reconstituted nasal epithelia maintained in air/liquid interphase conditions with 100 nM and 300 nM ABBV-744 and performed SARS-CoV-2 or mock infections (Fig. 6a). First, we found that ABBV-744 treatment reduced *ACE2* levels in these conditions (Fig. 6b). Apical supernatants did not show significant changes in viral RNA concentrations at two or four days post-infection (Extended Data Figure 7). Intracellular viral RNA concentrations, however, were significantly decreased in the ABBV-744 conditions (Fig. 6c). Furthermore, epithelial barrier integrity, as measured by trans-epithelial electrical resistance (Fig. 6d), and cytotoxicity (Fig. 6e), were rescued in infected cells treated with ABBV-744. Thus, ABBV-744 partially inhibited SARS-CoV-2 replication and fully rescued epithelial barrier integrity in a primary human nasal epithelial model.

**Figure 6:**
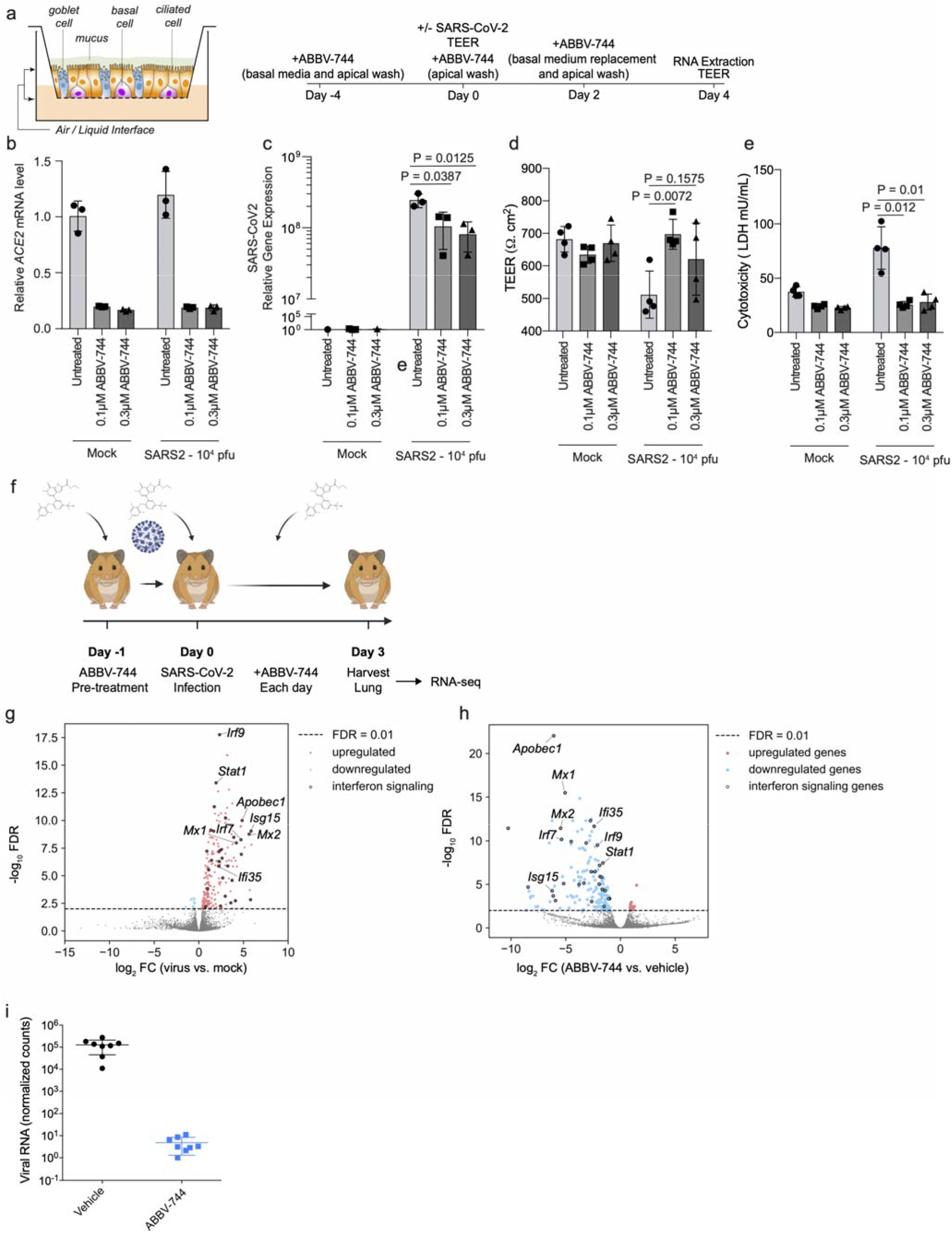
Brd2 inhibitors prevent cytotoxicity and reduce SARS-CoV-2 infection in human primary nasal epithelia and inhibit SARS-CoV-2 infection in Syrian Hamsters. **a,** Experimental design for experiment in reconstructed human nasal epithelia. **b**, ACE2 transcript levels relative to the average of GAPDH, TFRC, RPL13, and ACTB as a function of ABBV-744 concentration and/or SARS-CoV-2 infection (right). **c,** intracellular SARS-CoV-2 gene expression (N) relative to the average of GAPDH, TFRC, RPL13, and ACTB as a function of ABBV-744 concentration and/or SARS-CoV-2 infection (right). P-value was determined using Student’s unpaired two tailed t-test. **d,** Transepithelial electrical resistance (TEER), a measure of epithelial barrier integrity, evaluated as a function of ABBV-744 concentration and/or SARS-CoV-2 infection. P-value was determined using Student’s unpaired two tailed t-test. **e,** Cytotoxicity, as measured by lactate dehydrogenase (LDH) release, evaluated as a function of ABBV-744 concentration and/or SARS-CoV-2 infection. P-value was determined using Student’s unpaired two tailed t-test. Experiments a-d are done in at least biological triplicates with error bars representing the standard deviation. **f,** Experimental design for Syrian hamster experiments. **g**, Volcano plot showing differentially expressed genes for Syrian hamsters lungs infected or not infected with SARS-CoV-2. A subset of ISGs for which BRD2 peaks were identified by CUT&RUN are labeled. **h**, Volcano plot showing differentially expressed genes for Syrian hamsters lungs infected with SARS-CoV-2 and treated with vehicle or ABBV-744 at 100nM. A subset of ISGs for which BRD2 peaks were identified by CUT&RUN are labeled. **i,** Normalized viral RNA counts for Syrian hamsters infected with SARS-CoV-2 and treated with vehicle or ABBV-744 at 100 nM.

Next, we tested if ABBV-744 could reduce SARS-CoV-2 infection in golden Syrian hamsters. Syrian hamsters provide a physiologically relevant model for SARS-CoV-2 infection, with high viral replication and signs of lung involvement ^38–41^. After 24-hour treatment with ABBV-744 or vehicle, hamsters were infected with SARS-CoV-2 (Fig. 6f) and treated daily with ABBV-744 or vehicle. Three days post-infection, the lungs of hamsters were harvested and subjected to RNA-seq. Infected, but untreated, hamsters showed marked up-regulation of a number of genes including ISGs when compared to uninfected controls (Fig. 6g). In contrast, infected hamsters treated with ABBV-744 showed a down-regulation of ISG (Fig. 6h) levels relative to vehicle-treated infected hamsters, confirming ABBV-744 activity. Remarkably, viral RNA counts were reduced by about five orders of magnitude in the ABBV-744 treated hamsters versus those treated with vehicle controls (Fig. 6i). Thus, Brd2 inhibition can dramatically decrease SARS-CoV-2 infection in Syrian hamsters.

## Discussion

Here, we demonstrate that Brd2 is necessary for *ACE2* expression in a number of different SARS-CoV-2 relevant systems. We also found that treatment with ABBV-744, a bromodomain inhibitor, can reduce SARS-CoV-2 viral RNA concentrations in primary human nasal epithelial cells and Syrian hamsters. These findings suggest that pharmacological BRD2 inhibitors may be of therapeutic benefit to prevent or reduce the impact of SARS-CoV-2 infection.

Our data suggest that Brd2 is an indirect regulator of *ACE2* transcription in COVID-19-relevant cell types. Our data show that Brd2 is required for interferon-mediated stimulation of *ACE2* expression, as both exogenous interferon stimulation and basal interferon stimulation of *ACE2* expression is blocked upon *BRD2* knockdown or pharmacological inhibition (Fig. 6i). This does not, however, preclude a more direct, and interferon-independent, regulatory mode. Our data also show that Brd2 activity is essential for the transcription of ISGs in cell culture and in Syrian hamsters. Based on our findings and the previous literature^36^, Brd2 regulation of ISG transcription is likely mediated by a reduction in Histone H2A.Z occupancy at these promoters. Taken together, this indicates that BRD2 could be a key regulator of the host response to SARS-CoV-2 infection.

The previously described^42^ interaction between the SARS-CoV-2 E protein and BRD2 might have evolved to manipulate gene expression during infection, including the expression of *ACE2*. In isolation, however, protein E overexpression in Calu-3 cells did not recapitulate expression changes resulting from BRD2 knockdown or inhibition (Fig 4a). These data suggest that there is no direct effect of Protein E on BRD2 function, or that other viral or host factors expressed during SARS-CoV-2 infection are required to modulate BRD2 function. Further studies are needed to define the function of the protein E-BRD2 interaction.

Several previous CRISPR screens aiming to uncover strategies to inhibit SARS-CoV-2 infection were carried out in cell lines in which an *ACE2* transgene was overexpressed^9,10^; these screens therefore failed to uncover *BRD2* as a regulator of endogenous *ACE2* expression. BRD2 did show a phenotype, however, in a CRISPR screen carried out in Vero-E6 cells (which express *ACE2* endogenously)^12^, although it was not further characterized in that study. These differences highlight the importance of conducting CRISPR-based screens in disease-relevant cell types.

There is a growing literature about the relationship between COVID-19 disease severity, *ACE2* expression, and interferon regulation^1–6^. Since ACE2 is known to promote recovery after lung injury and that SARS-CoV-2 manipulates the host interferon response^43–45^, the mis-regulation of these two pathways may play a major role in enhancing the severity of COVID-19. Our data suggest that Brd2 is central to this regulatory network and, therefore, pharmacological targeting of Brd2 may be a promising therapeutic strategy for the treatment of COVID-19: Brd2 inhibition could both block viral entry, through ACE2 downregulation, and act as an “emergency-brake” for mis-regulated patient immune responses to COVID-19, via down-regulation of ISGs.

## Methods

### Cell Culture

Calu-3 cells were cultured in RPMI 1640 (Life Technologies 22400-105) with 10% FBS (VWR 89510-186), 1% Pen/Strep (Life Technologies 15140122), and 5 mM Glutamine (Life Technologies 25030081) at 37 °C and 5% CO_2_ Cells were split by treating with TrypLE (Life Technologies 12604013) for 15 minutes, quenching with media and spun down at 200xg for 5 minutes. At Institut Pasteur, where virus infections were carried out, Calu-3 cells were cultured in MEM (Gibco 11095-080) with 20% FBS (Gibco A3160801), 1% Pen/Strep (Gibco 15140-122), 1% NEAA (Sigma-Aldrich M7145) and 1 mM Sodium pyruvate (Sigma-Aldrich S8636). They were split in Trypsin-EDTA 0.05% (Gibco 11580626).

HEK293 cell culture and production of lentivirus was performed as previously described^46^.

A vial of STR authenticated Caco-2 cells was obtained from the UCSF Cell and Genome Engineering Core (CGEC). Caco-2 cells were cultured in EMEM (ATCC, 30-2003) with 20% FBS (VWR 89510-186), 1% Pen/Strep (Life Technologies 15140122), and 5 mM Glutamine (Life Technologies 25030081) at 37 °C and 5% CO2.

A vial of A549 cells was obtained from Davide Ruggero’s lab as a gift. A549 cells were cultured in DMEM (Thermo Fisher Scientific, 10313-039) with 10% FBS (VWR 89510-186), 1% Pen/Strep (Life Technologies 15140122), and 5 mM Glutamine (Life Technologies 25030081) at 37 °C and 5% CO2.

Human iPSC-derived cardiomyocytes were generated and cultured as previously described^30^, from AICS90 iPSCs (Allen Institute Cell Catalog). Drugs were added on day 69 of differentiation, and cardiomyocytes were harvested for analysis on day 72.

Normal human bronchial epithelia (Mattek NHBE-CRY) were cultured following the supplier’s instructions.

### Generation of the Calu-3 ACE2 knockout line

The polyclonal ACE2 knockout Calu-3 cell line was generated using the Gene KO kit V2 from Synthego, using three sgRNAs targeting ACE2 with the following protospacer sequences sRNA1: 5’-GACAUUCUCUUCAGUAAUAU-3’, sgRNA2: 5’-AAACUUGUCCAAAAAUGUCU-3’ and sgRNA3: 5’-UUACAGCAACAAGGCUGAGA-3’. Single guide RNAs (sgRNAs) were designed according to Synthego’s multiguide gene knockout kit^47^. Briefly, two or three sgRNAs are bioinformatically designed to work in a cooperative manner to generate small, knockout causing, fragment deletions in early exons. These fragment deletions are larger than standard indels generated from single guides. The genomic repair patterns from a multiguide approach are highly predictable on the basis of the guide spacing and design constraints to limit off-targets, resulting in a higher probability protein knockout phenotype.

The ribonucleoprotein (RNP) complex with a ratio of 4.5 to 1 between sgRNA and Cas9 was delivered following the protocol of the SE Cell Line 4D-NucleofectorTM X Kit (Lonza, V4XC-1012), using the nucleofection program DS-130 on the Lonza 4D X unit. 72 hours post transfection, genomic DNA was extracted to serve as the template for PCR amplification of the region that covers the sites targeted by the sgRNAs with the following two primers: ACE2-F: 5’-CTGGGACTCCAAAATCAGGGA-3’ and ACE2-R: 5’-CGCCCAACCCAAGTTCAAAG-3’. Sanger sequencing reactions using the sequencing primer ACE2-seq: 5’-CAAAATCAGGGATATGGAGGCAAACATC-3’ were then performed, and the knockout efficiency was determined to be 80% via ICE software from Synthego^48^ (https://ice.synthego.com/#/).

### Generation of the Calu-3 CRISPRi line

The parental Calu-3 line was obtained from the UCSF Cell and Genome Engineering Core. Calu-3 cells were cultured at 37 °C with 5% CO2 in EMEM media containing 10% FBS, 100 units/ml streptomycin, 100 μg/ml penicillin, and 2 mM glutamine. To generate the CRISPRi lines, ~3×10^6^ cells were seeded into media containing lentiviral particles packaging dCas9-BFP-KRAB under a UCOE-SFFV promoter^49^. Five days post infection, BFP-positive cells were sorted using a BD Fusion. To validate the CRISPRi line, Calu-3-CRISPRi cells were transduced with lentiviral particles expressing non-targeting sgRNA (protospacer 5’-GCTCCCAGTCGGCACCACAG-3’) or *CD81*-targeting sgRNA (protospacer 5’-GGCCTGGCAGGATGCGCGG-3’). CD81 expression was measured 7 days post-transduction by dislodging cells with TrypLE and live cells were stained with APC-conjugated anti-human CD81 antibody (Biolegend 349509). CD81 expression was assessed on a BD LSRII with >90% of Calu3-CRISPRi cells with CD81 knocked down compared to a non-targeting sgRNA control.

### Spike-RBD binding assay

Recombinant biotinylated SARS-CoV-2 spike Spike-receptor-binding domain with a C-terminal human IgG Fc domain fusion (referred to as Spike-RBD) was prepared as previously described^50^. Calu-3 cells were grown in 96-well flat bottom plates until >50% confluent. Media was aspirated and cells were washed once with PBS. Cells were then treated with TrypLE to release them from the plate, RPMI 1640 media was added to dilute TrypLE, and cells were pelleted by centrifugation at 200xg for five minutes. From this point on, all steps were carried out on ice. Cells were incubated in 3% BSA (Sigma Aldrich A7030) in DPBS (Sigma-Aldrich D8537) for 15 minutes to block and washed twice in 3% BSA in DPBS by centrifugation at 200xg for five minutes in v-bottom plates, followed by resuspension. Spike-RBD was diluted in 3% BSA to appropriate concentrations and incubated with cells for 30 minutes on ice. Cells were then washed twice with 3% BSA in DPBS and incubated with Anti-Strep PE-Cy7 (Thermofisher SA1012) at 5 μg/mL. Cells were washed twice and subjected to flow cytometry on a FACS Celesta in HTS mode. Cells were gated to exclude doublets and the median PE-Cy7 signal was calculated for each sample. The gating strategy is shown in Supplemental Figure 2. EC_50_ values and their 95% confidence intervals were calculated by fitting the RBD binding data into a Sigmoidal, 4PL model in Prism 6.

### CRISPRi Screen

Calu-3 cells were infected with the H1 CRISPRi sgRNA library^22^ as described^46^ and selected using treatment with 1 μg/mL puromycin for 3 days. After selection, cells were stained with 10 nM Spike-RBD as described above or for TFRC as previously described^46^ and subjected to FACS, where cells were sorted into top 30% and bottom 30% based on high and low expression of TFRC or Spike-RBD. Because of viability and stickiness known for Calu-3 cells, coverage was lower than optimal, at 200-fold over the library diversity. Sorted populations were spun down at 200xg for five minutes and genomic DNA was isolated as described^46^. sgRNA cassettes were amplified by PCR and sequencing and analysis was performed as described^46^ but with an FDR of 0.1 rather than 0.05 or 0.01 due to noise.

### Validation of screening hits

Individual sgRNAs were selected based on phenotypes in the primary screens and cloned into a lentiviral expression vector as described^46^. Protospacer sequences of these sgRNAs are provided in **Extended Data Table 5**. Cells expressing sgRNAs were selected using treatment with 1 μg/mL puromycin for 3-7 days.

### Drug treatments

Drugs (ABBV-744 Selleckchem S8723, JQ1 - Sigma Aldrich SML1524, dBET6 - Selleckchem S8762) were dissolved in DMSO or water as per manufacturer’s instructions. Cells were treated with drugs for 72 hours with media changes performed every 24 hours with media containing fresh drug.

### Interferon Treatments

Interferon Beta (R&D systems 8499-IF) was dissolved per the manufacturer’s instructions. Cells were treated with IFNβ for 72 hours with media changes performed every 24 hours with media containing fresh IFNβ.

### qPCR

qPCR was performed and analyzed as described^46^. Primers: *ACE2* forward: GGTCTTCTGTCACCCGATTT; *ACE2* reverse: CATCCACCTCCACTTCTCTAAC; *ACTB* forward: ACCTTCTACAATGAGCTGCG; *ACTB* reverse: CCTGGATAGCAACGTACATGG; *IRF9* forward: GCCCTACAAGGTGTATCAGTTG; *IRF9* reverse: TGCTGTCGCTTTGATGGTACT; *IFNAR1* forward: AACAGGAGCGATGAGTCTGTC; *IFNAR1* reverse: TGCGAAATGGTGTAAATGAGTCA; *STAT1* forward: CAGCTTGACTCAAAATTCCTGGA; *STAT1* reverse: TGAAGATTACGCTTGCTTTTCCT.

### Western Blotting

Cells from one confluent well of a six-well plate were lysed in RIPA buffer plus c0mplete EDTA-free protease inhibitor tablets (Roche 11873580001) and spun for 10 minutes at 21,000xg at 4°C. The pellet was removed and a BCA assay (Thermofisher 23225) was performed on the remaining supernatant. Lysate volumes with equivalent protein content were diluted with SDS-PAGE loading dye and subjected to gel electrophoresis on 4-12% BisTris SDS-PAGE gels (Life Technologies NP0322). Gels were then transferred and blocked in 5% NFDM for 1 hour at RT. Antibodies in fresh 5% NFDM were added (Mouse monoclonal GAPDH 1:10,000; Goat polyclonal ACE2 [R&D Tech AF933] 1:200; Rabbit monoclonal BRD2 [abcam 197865] 1:5,000) and incubated at 4 °C for at least 16 hours. Membranes were washed 4x with TBS + 0.1% Tween-20 and incubated with secondary antibodies (1:10,000 Donkey Anti-Goat-800 [LICOR 926-32214], LICOR Donkey Anti-Mouse-680 [LICOR 926-68072], HRP Donkey anti-rabbit [CST 7074P2]). Membranes were visualized using a LiCOR or Femto HRP kit (Thermofisher 34094). Uncropped images of Western blots are provided as Supplemental Figure 1.

### Virus

The SARS-CoV-2 strain used (BetaCoV/France/IDF0372/2020 strain) was propagated once in Vero-E6 cells and is a kind gift from the National Reference Centre for Respiratory Viruses at Institut Pasteur, Paris, originally supplied through the European Virus Archive goes Global platform.

### Cytotoxicity measurements of Calu3 cells

30,000 Calu-3 cells per well were seeded into Greiner 96-well white bottom plates and incubated for 48 hours at 37°C, 5% CO_2_. Then, cells were treated with identical drug concentrations as in the infection assays for 5 days by refreshing the media with 100μL per well fresh drug-containing media every 24 hours. Cell viability was then assayed by adding 100μL per well of CellTiter-Glo 2.0 (Promega) and incubated for 10 minutes at room temperature. Luminescence was recorded with an Infinite 200 Pro plate reader (Tecan) using an integration time of 1s.

### Virus infection assays

30,000 Calu-3 cells per well were seeded into 96-well plates and incubated for 48 hours at 37°C, 5% CO_2_. At the time of infection, the media was replaced with virus inoculum (MOI 0.1 PFU/cell) and incubated for one hour at 37°C, 5% CO_2_. Following the one-hour adsorption period, the inoculum was removed, replaced with fresh media, and cells incubated at 37°C, 5% CO_2_. 24h, 48h and 72h post infection, the cell culture supernatant was harvested, and viral load assessed by RT-qPCR as described previously^42^. Briefly, the cell culture supernatant was collected, heat inactivated at 95°C for 5 minutes and used for RT-qPCR analysis. SARS-CoV-2 specific primers targeting the N gene region: 5′-TAATCAGACAAGGAACTGATTA-3′ (Forward) and 5′-CGAAGGTGTGACTTCCATG-3′ (Reverse) were used with the Luna Universal One-Step RT-qPCR Kit (New England Biolabs) in an Applied Biosystems QuantStudio 6 thermocycler or an Applied Biosystems StepOnePlus system, with the following cycling conditions: 55°C for 10 min, 95°C for 1 minute, and 40 cycles of 95°C for 10 seconds, followed by 60°C for 1 minute. The number of viral genomes is expressed as PFU equivalents/mL, and was calculated by performing a standard curve with RNA derived from a viral stock with a known viral titer.

### Plaque assays

Viruses were quantified by plaque-forming assays. For this, Vero E6 cells were seeded in 24-well plates at a concentration of 1 × 10□ cells per well. The following day, tenfold serial dilutions of individual virus samples in serum-free DMEM medium were added to infect the cells at 37□°C for 1 h. After the adsorption time, a solid agarose overlay (DMEM, 10% (v/v) PBS and 0.8% agarose) was added. The cells were incubated for a further 3 days prior to fixation with 4% formalin and visualization using crystal violet solution.

### CUT&RUN

*CUT&RUN* was performed with 1 million Calu-3 cells. Cells were removed from the plate by treatment with Versene (Life Technologies 15040066) for 20 minutes and resuspended in fresh media. They were spun down and washed twice with DPBS before proceeding with the CUTANA *CUT&RUN* kit (Epicypher 14-0050). The experiment was performed with the included IgG and H3K4Me control antibodies and the BRD2 antibody (abcam 197865) as well as *E.coli* spike-in DNA according to the kit protocol.

### QuantSeq analysis

Raw sequencing reads from QuantSeq were trimmed using Trimmomatic^51^ (v0.39, PMID: 24695404) and mapped to the human reference transcriptome (GRCh38, GENCODE Release 36) using Salmon^52^ (v1.3.0) to obtain transcript abundance counts. Gene-level count estimates were obtained using tximport^53^ (v1.18.0) with default settings. Subsequently, differential gene-expression analyses were performed using the glmQLFTest method implemented in the edgeR package^54^ (v3.28.1). Cluster^55^ (v3.0) was used for hierarchical clustering and Java TreeView^56^ (v1.1.6r4) for visualization.

### CUT&RUN analysis

CUT&RUN analysis was performed as previously described^57^. Briefly, paired-end reads were mapped to the human genome GRCh38 using Bowtie2 (v2.3.4.1) with options: --end-to-end --very-sensitive --no-unal --no-mixed --no-discordant --phred33 -I 10 -X 1000. Sparse Enrichment Analysis for CUT&RUN (SEACR^58^, https://seacr.fredhutch.org/) was used for peak calling. H3K4me3 and BRD2 peaks were normalized to IgG control. Published BRD2 ChIP-seq data in human lung cells was obtained from ChIP-Atlas (https://chip-atlas.org/). The Integrative Genomics Viewer (IGV, igv.org) was used for visualization.

#### SARS-CoV-2 infection of reconstructed human nasal epithelia

MucilAir™, corresponding to reconstructed human nasal epithelium cultures differentiated in vitro for at least 4 weeks, were purchased from Epithelix (Saint-Julien-en-Genevois, France). The cultures were generated from pooled nasal tissues obtained from 14 human adult donors. Cultures were maintained in air/liquid interface (ALI) conditions in transwells with 700 μL of MucilAir™ medium (Epithelix) in the basal compartment, and kept at 37°C under a 5% CO2 atmosphere.

SARS-CoV-2 infection was performed as previously described^37^. Briefly, the apical side of ALI cultures was washed 20 min at 37°C in Mucilair™ medium (+/− drug) to remove mucus. Cells were then incubated with 10^4^ plaque-forming units (pfu) of the isolate BetaCoV/France/IDF00372/2020 (EVAg collection, Ref-SKU: 014V-03890; kindly provided by S. Van der Werf). The viral input was diluted in DMEM medium (+/− drug) to a final volume 100 μL, and left on the apical side for 4 h at 37°C. Control wells were mock-treated with DMEM medium (Gibco) for the same duration. Viral inputs were removed by washing twice with 200 μL of PBS (5 min at 37°C) and once with 200 μL Mucilair™ medium (20 min at 37°C). The basal medium was replaced every 2-3 days. Apical supernatants were harvested every 2-3 days by adding 200 μL of Mucilair™ medium on the apical side, with an incubation of 20 min at 37°C prior to collection.

For ABBV-774 treatment, cultures were pretreated for 4 days with 100 nM or 300 nM of the drug. For this pretreatment, ABBV-744 was added to an apical wash at day −4, and to the basal compartment from day −4 to day 0. The drug was then added on the apical side during viral adsorption at day 0, and then every 2-3 days to both the apical wash and the basal compartment throughout the infection.

### Transepithelial electrical resistance (TEER) measurement

The apical side of transwell cultures was washed for 20 min at 37°C in Mucilair™ medium. Transwell were then transferred in a new 24-well plate and DMEM medium was added to both the apical (200 μL) and basal (700 μL) sides. The TEER was then measured using an Evom3 ohmmeter (World Precision Instruments).

### LDH cytotoxicity assay

Diluted culture supernatants (1:25) were pre-treated with Triton-X100 1% for 2 h at RT for viral inactivation. Lactate dehydrogenase (LDH) dosage was performed using the LDH-Glo™ Cytotoxicity Assay kit (Promega) following manufacturer’s instructions. Luminescence was measured using an EnSpire luminometer (Perkin Elmer).

### Viral RNA quantification

Apical supernatants were stored at −80°C until thawing and were diluted 4-fold in PBS for quantification in a 96-well PCR plate. Supernatants were then inactivated for 20 min at 80°C. One μL of supernatant was directly added to 4μL of PCR reaction mix for SARS-CoV-2 RNA quantification. PCR was carried out in a final volume of 5 μL per reaction in 384-well plates using the Luna Universal Probe One-Step RT-qPCR Kit (New England Biolabs) with SARS-CoV-2 N-specific primers (Forward 5′-TAA TCA GAC AAG GAA CTG ATT A-3′; Reverse 5′-CGA AGG TGT GAC TTC CAT G-3′) on a QuantStudio 6 Flex thermocycler (Applied Biosystems). Standard curve was established in parallel using purified SARS-CoV-2 viral RNA.

### Tissue RNA quantification

ACE2 and SARS-CoV-2 expression was quantified in epithelial cells by real-time quantitative PCR. The epithelial cultures were washed in ice cold PBS and then lyzed in 150 μL of Trizol reagent (Thermofisher Scientific) added to the apical side of the insert for 5 min. RNA was purified using the Direct-zol miniprep kit (ZR2080, Zymo Research). Transcripts of genes of interest (*ACE2*, SARS-CoV-2 N gene) were amplified in a final volume of 5 μL per reaction in 384-well plates using the Luna Universal Probe One-Step RT-qPCR Kit (New England Biolabs) on a QuantStudio 6 Flex thermocycler. RT-qPCR results were normalized to the mean expression of 4 reference genes (*GAPDH, TFRC, ALAS1, RLP13*) to compute relative gene expression, as described previously^37^. The *ACE2* primers used were ACE2-For 5’-TGG GAC TCT GCC ATT TAC TTA C-3’ and ACE2-Rev 5′-CCA GAG CCT CTC ATT GTA GTC T-3.

#### *In vivo* infections

All animal infections were conducted at the Icahn School of Medicine at Mount Sinai, under biosecurity level 3 (BSL-3) facility of the Global Health and Emerging Pathogen Institute approved by the Institutional Animal Care and Use Committee at Icahn School of Medicine at Mount Sinai under protocol number IACUC#20-0743. 6- to 8-week-old male Golden Syrian hamsters (*Mesocricetus auratus*) were purchased from Jackson Laboratories, housed in pairs, and fed *ad libitum*. On the day of infection, animals were anesthetized by administration of 100 ul of a ketamine HCl/xylazine (4:1) mix by intra-peritoneal injection and infected intranasally with 1000 plaque forming units of SARS-CoV-2 USA-WA1/2020 diluted in 100 ul of PBS. At indicated time points, animals were treated with 1 mL of ABBV-744 by oral gavage. The drug was prepared fresh daily to a final concentration of 20 mg/kg in 0.5% HPMC /0.5% Tween 80 in water. On day 3 after infection animals received 100 ul of a mix of pentobarbital/PBS (1:4) intraperitoneally, and once anesthetized they were cervically dislocated and lung lobes collected in 1 mL of Trizol reagent. Tissues were homogenized in a Tissue lyser for 2 cycles of 40 seconds, spun down for 5 minutes at 8000g and supernatants stored at −80C for plaque assay or RNA extraction.

### RNA extraction

RNA extraction was performed following instructions from the manufacturer of TRIzol reagent (Invitrogen). Briefly, 1/5 volume of chloroform was added to the lung supernatants in TRIzol, phases were separated by centrifugation and RNA was precipitated by overnight incubation with isopropanol at −20C. The RNA pellet was washed with ethanol 70% and resuspended in RNase-free water. RNA was quantified by nanodrop and resuspended to a final concentration of 100 ng/ul in water.

### RNA-Seq

1 ug of RNA was used as starting material for library preparation. The kit employed was TruSeq RNA Library Prep Kit v2 (Illumina) over polyadenylated RNA and the manufacturer’s instructions were followed. The sequencing was performed on an Illumina NextSeq 500 instrument. The raw reads obtained from the run were aligned against the Syrian golden hamster genome (MesAur1.0) in the Basespace platform by Illumina, with the tool “RNA-Seq Alignment”.

## Acknowledgements

We thank members of the Kampmann, Vignuzzi and Conklin labs, as well as Vijay Ramani, Davide Ruggero, Melanie Ott and her lab and other members of the UCSF QBI Coronavirus Research Group (QCRG) for helpful discussions. We thank Kun Leng for feedback on the manuscript. We thank the Gladstone Stem Cell Core for help with cardiomyocyte production. AJS is supported by NIH F32AG063487. GNR is supported by the NSF Graduate Research Fellowship Program and UCSF Discovery Fellowship. SAL was a Merck Fellow of the Helen Hay Whitney Foundation. IL was supported by an NSF GRFP. MK is a Chan Zuckerberg Biohub Investigator.

## Author Contributions

RT, AJS and MK conceptualized the overall project, analyzed results and prepared the manuscript, with input from all co-authors. VR, AMK and QDT performed and analyzed live-virus experiments in Calu-3 cells with guidance from MV. RR performed and analyzed human nasal epithelia experiments with guidance from LAC. LC performed and analyzed Syrian hamster experiments with guidance from BRT. GNR and SJR performed and analyzed experiments with cardiomyocytes, with guidance from BC. RT, AJS, MC and XG performed and analyzed all other experiments, with guidance from MK. JW performed and analyzed basal interferon signaling knockdown experiments in Calu-3 cells with guidance from RT. NL analyzed QuantSeq data with guidance from RT. SL, IL and JAW generated Spike-RBD; JN and JSW generated the Calu-3 CRISPRi cell line JCS, JO, TM and KH designed and provided sgRNAs to generate the ACE2 KO cell line.

## Data availability statement

Source data for immunoblots are provided in Supplementary Fig. 1. Gating strategies for flow cytometry experiments are provided in Supplementary Fig. 2. Sequencing data are provided available on NCBI Gene Expression Omnibus (GEO) with the following accession numbers: GSE165025 (RNA sequencing data associated with Fig. 4), GSE182993 (CUT&RUN data associated with Fig. 5), and GSE182994 (RNA sequencing data associated with Fig. 6f–h. There are no restrictions on data availability.

## Code availability statement

Analysis of CRISPRi screen results was carried out using custom code (MAGeCK-iNC) developed in the Kampmann lab, which was previously described^46^ and is freely available at https://kampmannlab.ucsf.edu/mageck-inc.

---

**Extended Data Figure 1:**
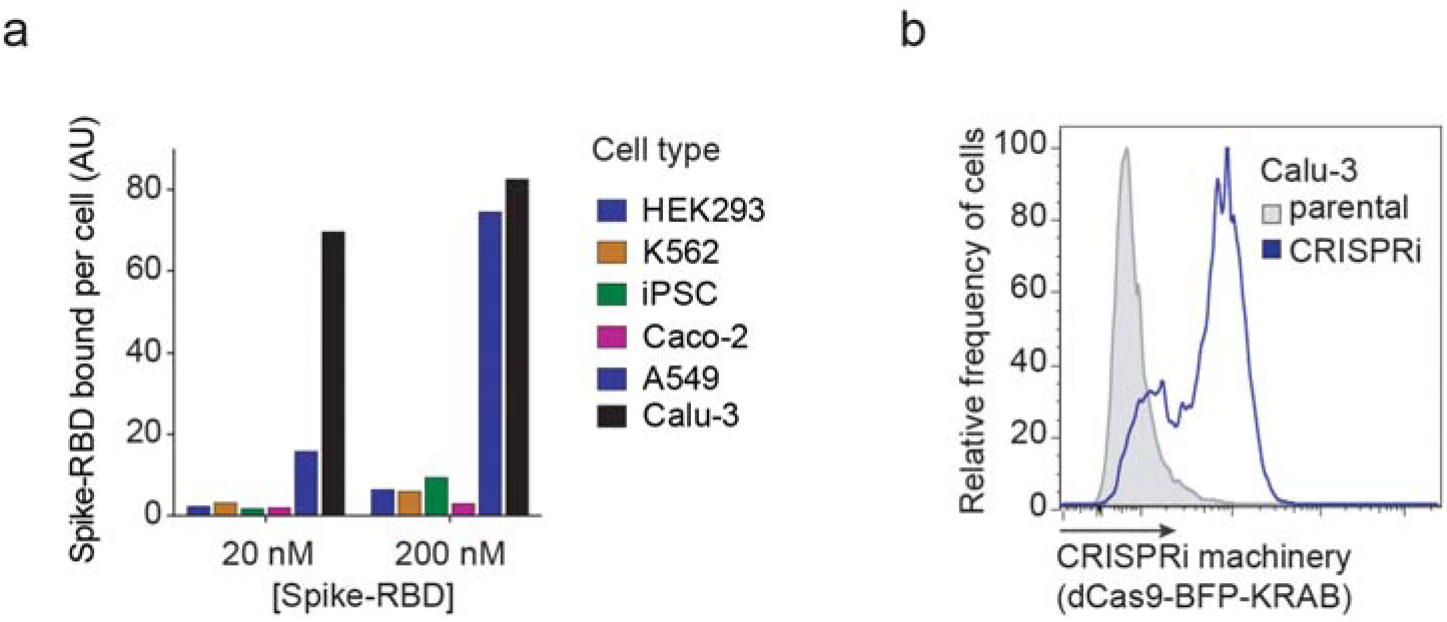
Calu-3 cells bind Spike-RBD specifically and were engineered to express CRISPRi machinery. **a**, Spike-RBD binding in different cell types at 20 nM and 200 nM Spike-RBD was quantified by flow cytometry. **b**, Expression of CRISPRi machinery (dCas9-BFP-KRAB) in the CRISPRi Calu-3 line indicated by the expression of BFP by flow cytometry.

**Extended Data Figure 2:**
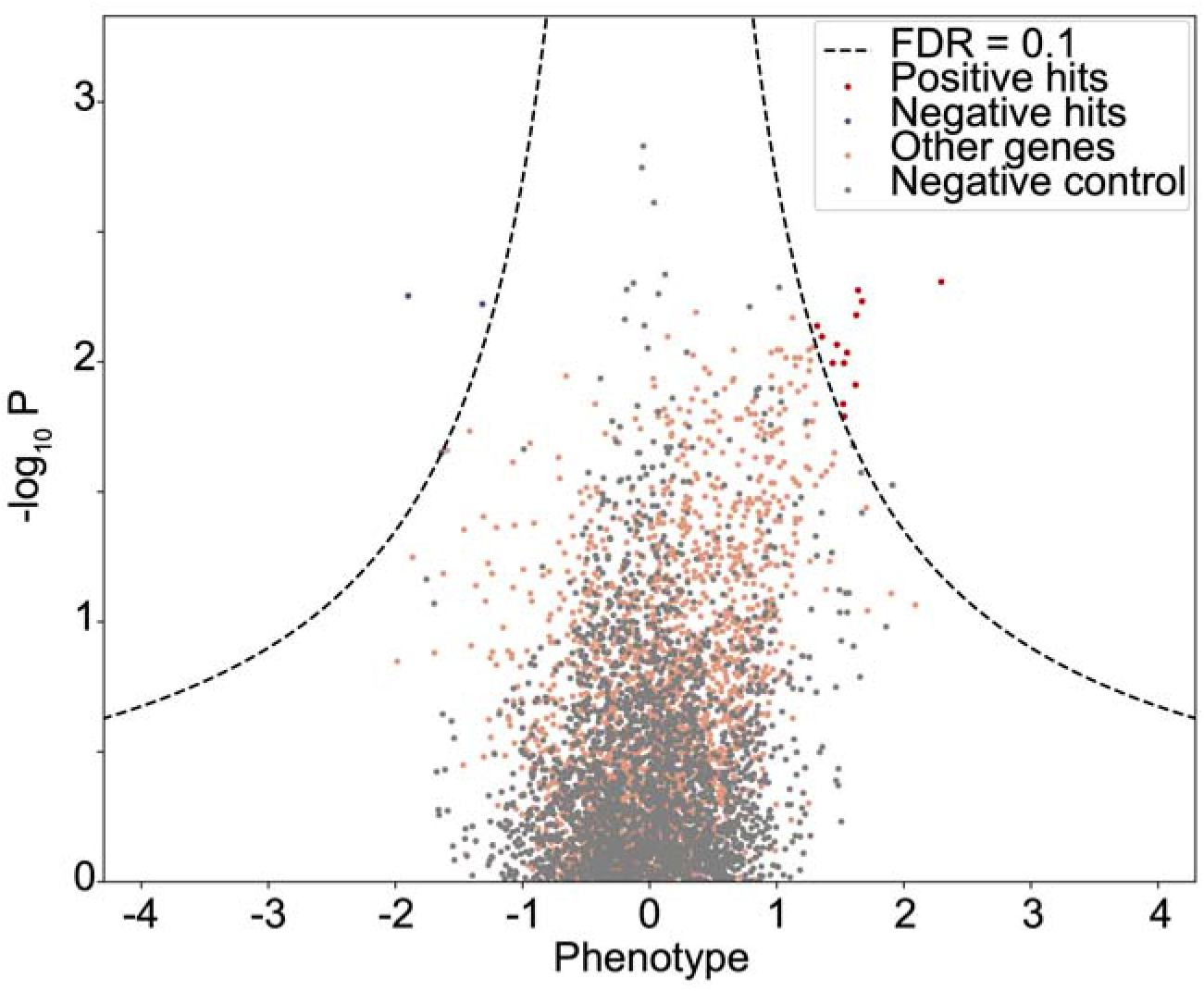
Volcano plot of Spike-RBD screen. Enrichment of sgRNAs targeting specific genes (colored dots) or non-targeting control sgRNAs plotted against the negative log of the P-value with a FDR of 0.1 shown (dashed lines).

**Extended Data Figure 3:**
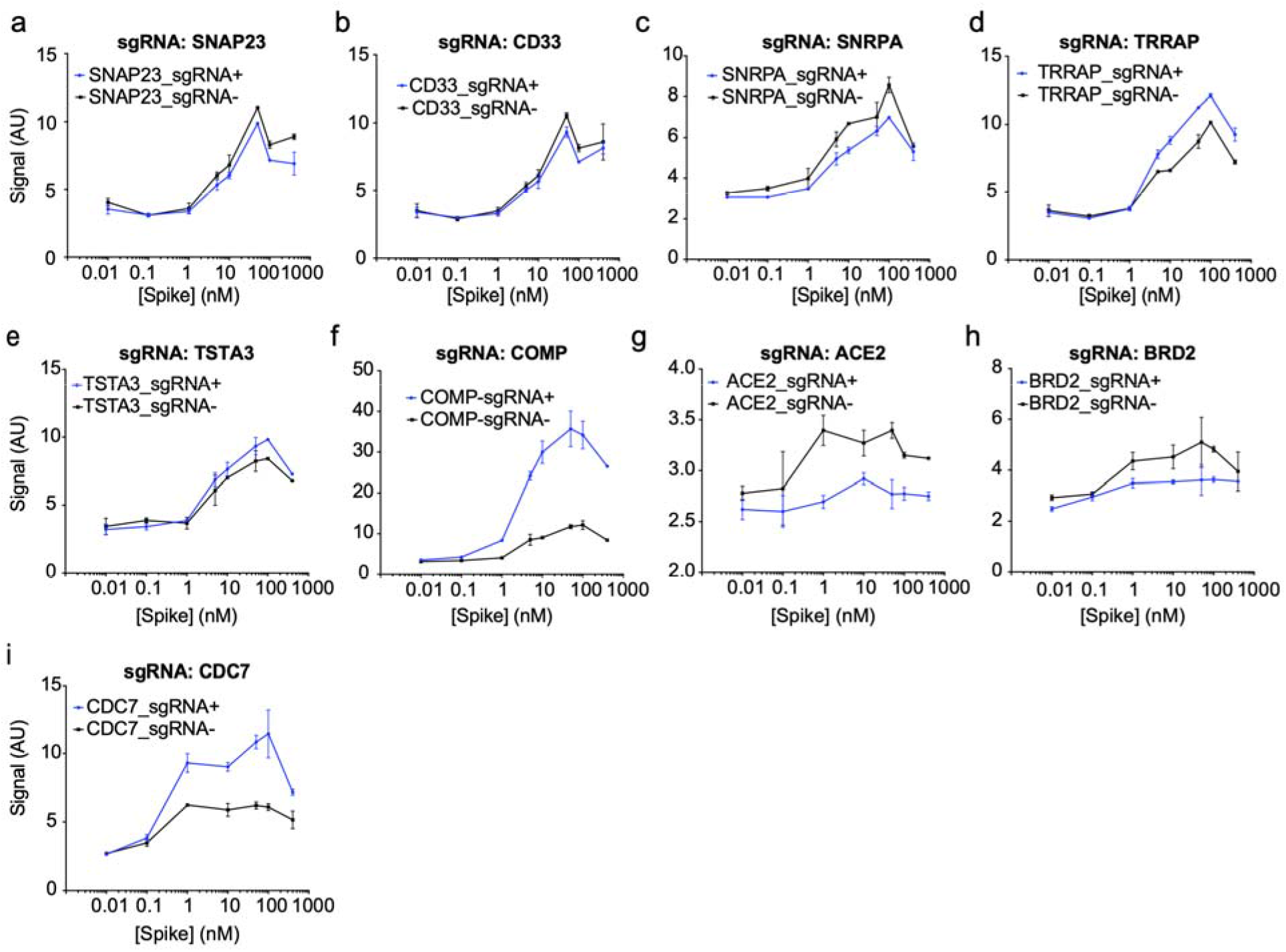
Individual sgRNA re-test of screening hits. **a-i,** Spike-RBD signal measured by flow-cytometry as a function of Spike-RBD concentration. Blue lines represent cells expressing the sgRNA targeting the gene of interest, black lines represent un-transduced control cells in the same well. Error bars represent s.d. from 2 independent wells.

**Extended Data Figure 4:**
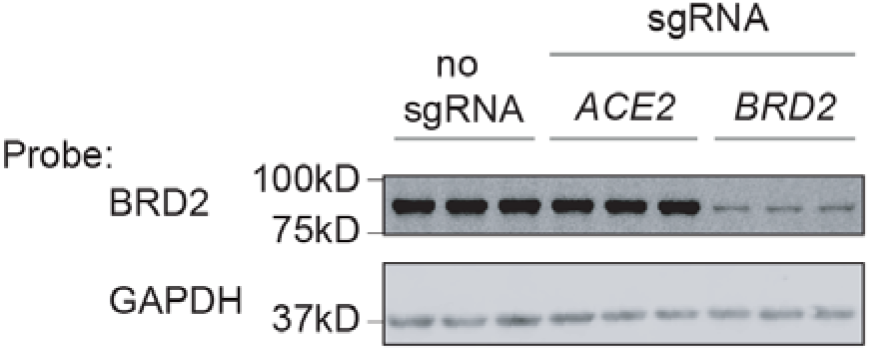
BRD2 is effectively knocked down by CRISPRi. Western blot for BRD2 and the loading control GAPDH in CRISPRi Calu-3 cells expressing no sgRNA or sgRNAs targeting ACE2 or BRD2. Three lanes represent samples from three independent wells.

**Extended Data Figure 5:**
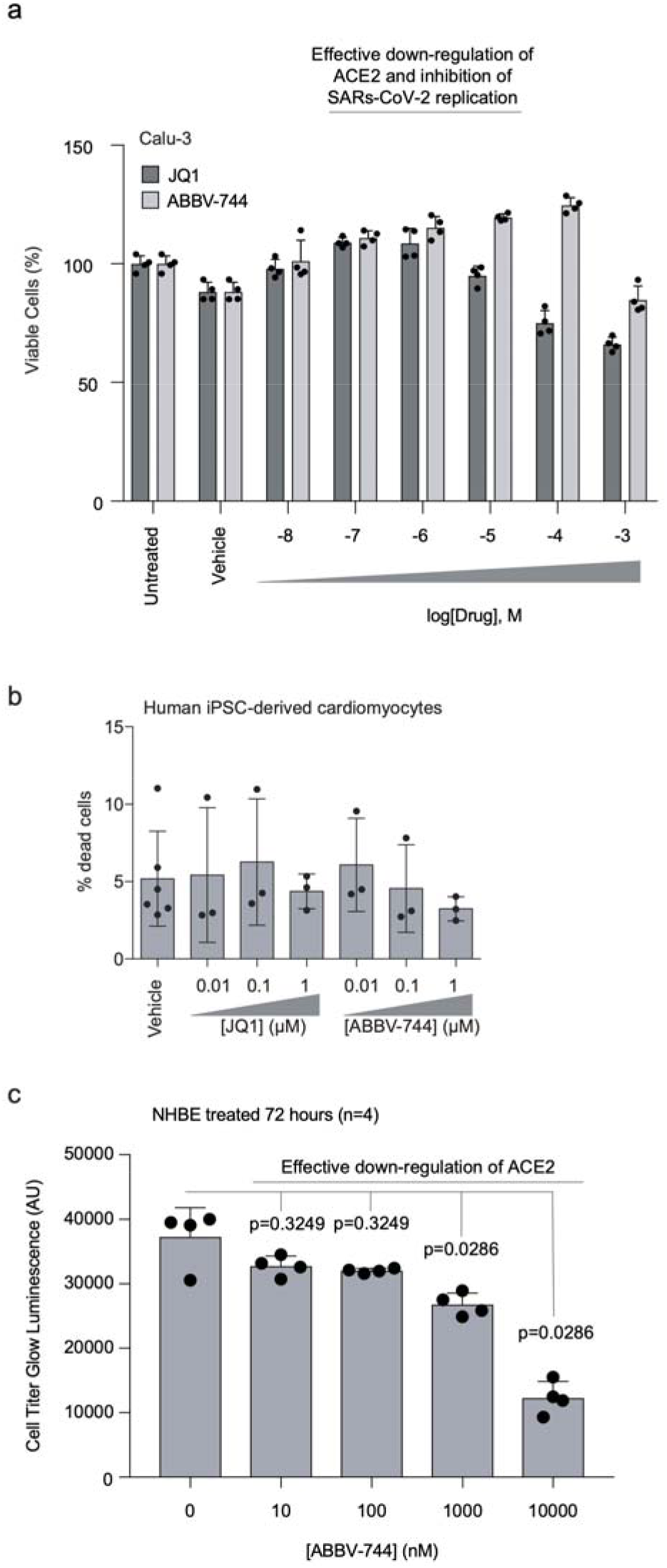
Non-toxic concentration range of BRD2 inhibitors. **a**, Calu-3 cells were treated with vehicle or the indicated concentrations of JQ1 or ABBV-744 for 5 days. Cell viability was then assayed with CellTiter-Glo 2.0 to calculate viability. Error bars represent the standard deviation of four biological replicates. **b**, Human iPSC-derived cardiomyocytes were treated for 72 hours with vehicle or the indicated concentrations of JQ1 or ABBV-744, and the percentage of dead cells was quantified as the ratio of propidium iodide-positive cells (dead cells) over Hoechst-positive cells (all cells). Error bars represent the standard deviation of three biological replicates (six biological replicates for the vehicle condition). **c,** Primary human bronchial epithelial (NHBE) cells were treated with ABBV-744 at the indicated concentrations for 72 hours and toxicity was assessed using CellTiter-Glo 2.0. Error bars represent the standard deviation of four biological replicates. P-values determined using Mann-Whitney two tailed test.

**Extended Data Figure 6:**
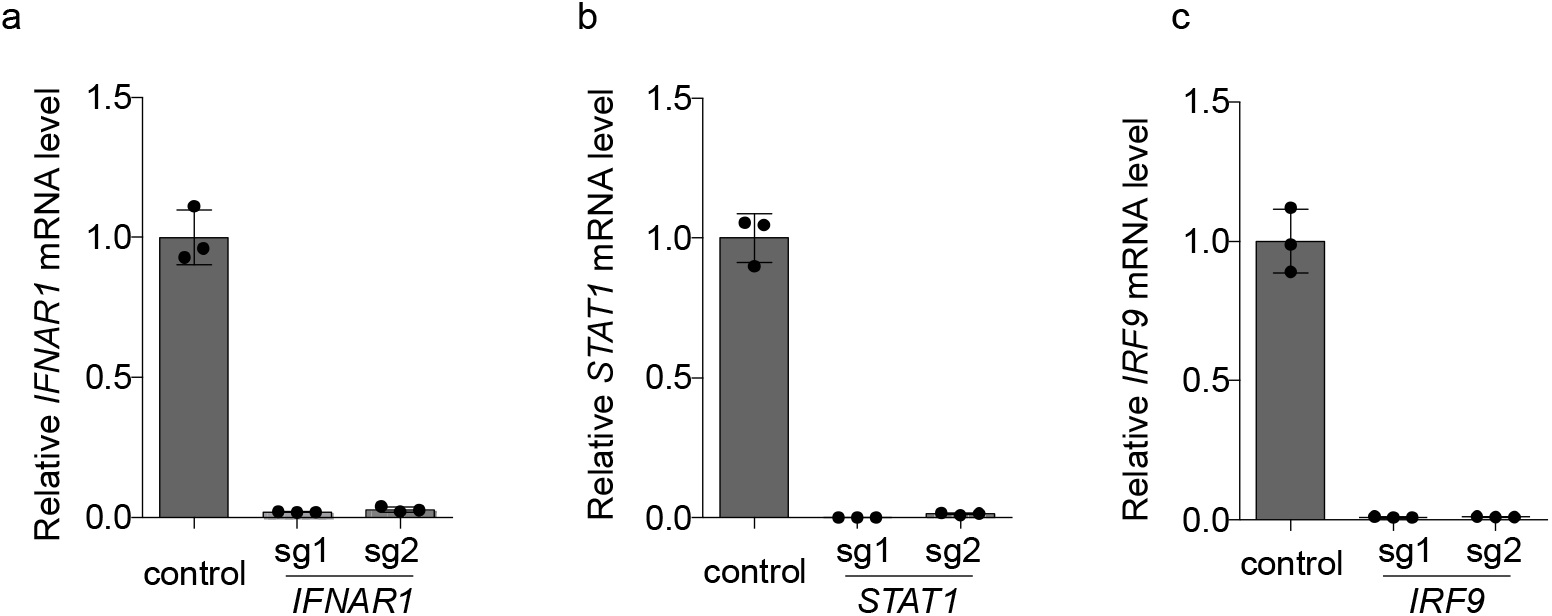
Validation of knockdown of interferon regulators by CRISPRi. **a-c,** Calu-3 cells expressing sgRNAs knocking down genes essential for interferon signal transduction assayed for transcript levels of sgRNA targets relative to *ACTB* by qPCR. Error is the standard deviation of three biological replicates.

**Extended Data Figure 7:**
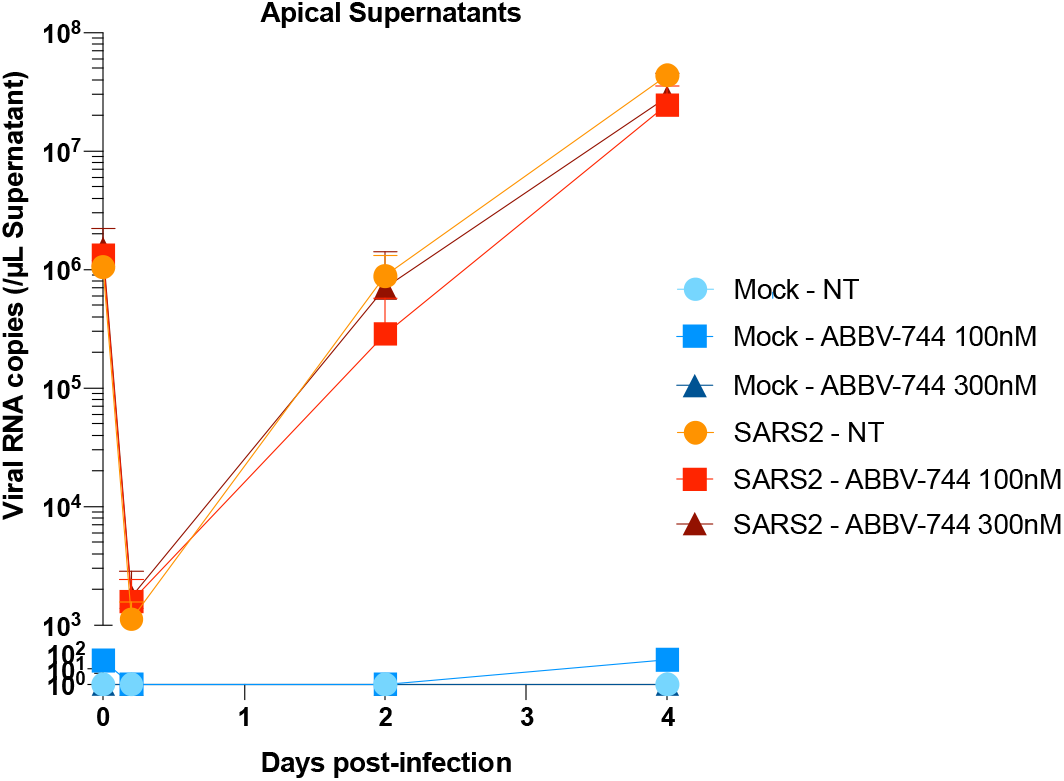
Viral replication in apical supernatants of reconstructed human nasal epithelia cultures. Apical supernatants of either infected or mock-infected nasal epithelia treated with ABBV-744 at the indicated concentrations or not treated (NT) were isolated and assayed for SARS-CoV-2 N RNA content. Experiments were done in biological quadruplicates with error bars representing the standard deviation.

## Extended Data Table Legends

**Extended Data Table 1: Phenotypes from CRISPRi screens for Spike-RBD and anti-TFRC binding.**

Results from CRISPRi screens for Spike-RBD and anti-TFRC binding were analyzed by the MAGeCK-iNC pipeline (see Methods for details) and are listed for all genes targeted by the H1 sgRNA library. Columns are: Targeted gene, targeted transcription start site, knockdown phenotype (epsilon), P value, and Gene score.

**Extended Data Table 2: Results from Quant-Seq experiments.**

The first six tabs show the results of differential gene expression analyses for *ACE2* knockdown, ABBV-744 treatment, *BRD2* knockdown, JQ1 treatment, SARS-CoV-2 protein E overexpression and *COMP* knockdown, respectively, using edgeR (see Methods for details). Columns are: Gene symbol, log_2_-fold change, log_2_ counts per million, F value, P value and FDR by the Benjamini-Hochberg method.

The ‘TPM’ tab shows the raw Transcripts Per Million (TPM) values for all samples. Columns: treatment conditions with 2 replicates each. Rows: all genes in the human transcriptome reference. The last tab provides the numerical values underlying the heatmap in Figure 4a. Columns: treatment conditions Rows: genes that are among top 50 differentially expressed genes in any of the conditions.

**Extended Data Table 3: Results from Syrian hamster RNA-seq.**

Results of differential gene expression analyses using edgeR for Syrian hamster lungs. First tab, SARS-CoV-2 infected compared to uninfected Syrian hamster lungs; Second tab, 100nm ABBV-744 compared to vehicle treated Syrian hamster lungs after SARS-CoV-2 infection. Columns are: Gene symbol, log_2_-fold change, log_2_ counts per million, F value, P value and FDR by the Benjamini-Hochberg method.

**Extended Data Table 4: Results from CUT&RUN experiments.**

BRD2 direct targets that are up- or down-regulated in the *BRD2* knockdown condition identified by the BETA analyses are listed. Columns are up-regulated targets and down-regulated targets.

**Extended Data Table 5: Protospacer sequences of individually tested sgRNAs.**

Protospacer sequences of individual sgRNAs used in Figure 1g are listed.

**Extended Data Table 6: All numerical data plotted in this paper**

All numerical data for each figure panel is contained in this table.

---

**Supplemental Figure 1:**
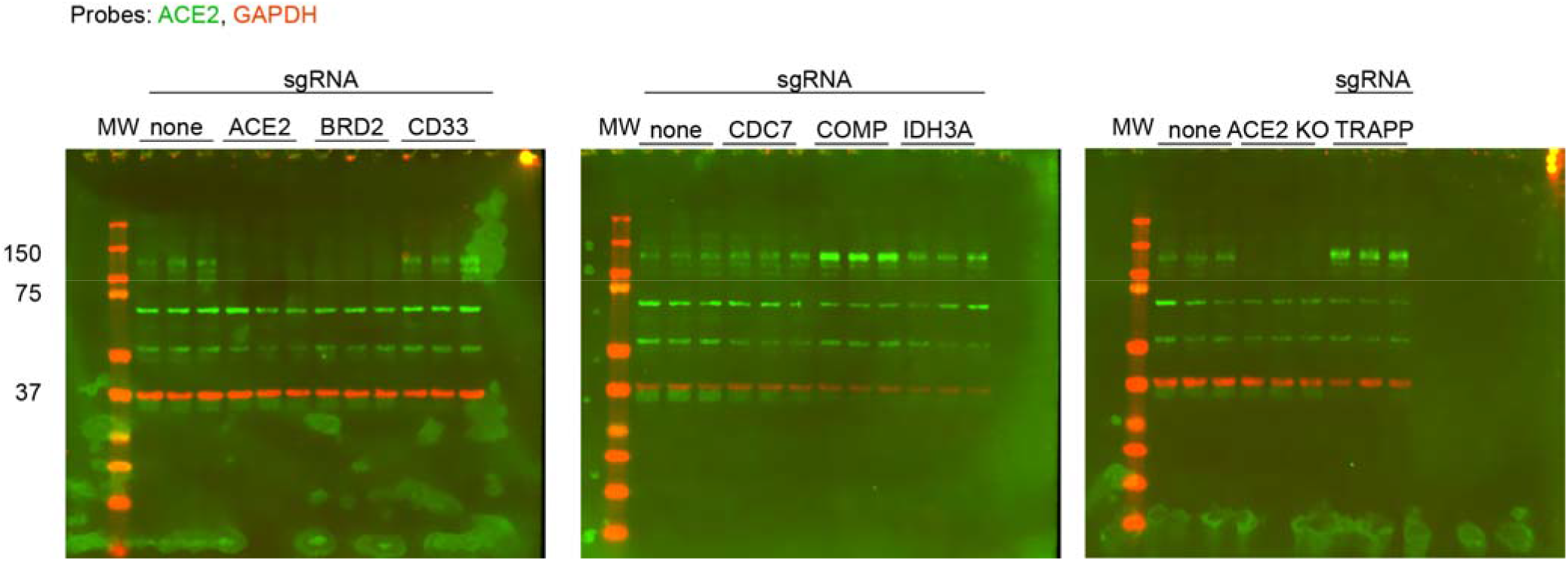
Full-size Western blots, associated with Fig. 2a and Extended Data Fig. 4.

**Supplemental Figure 2:**
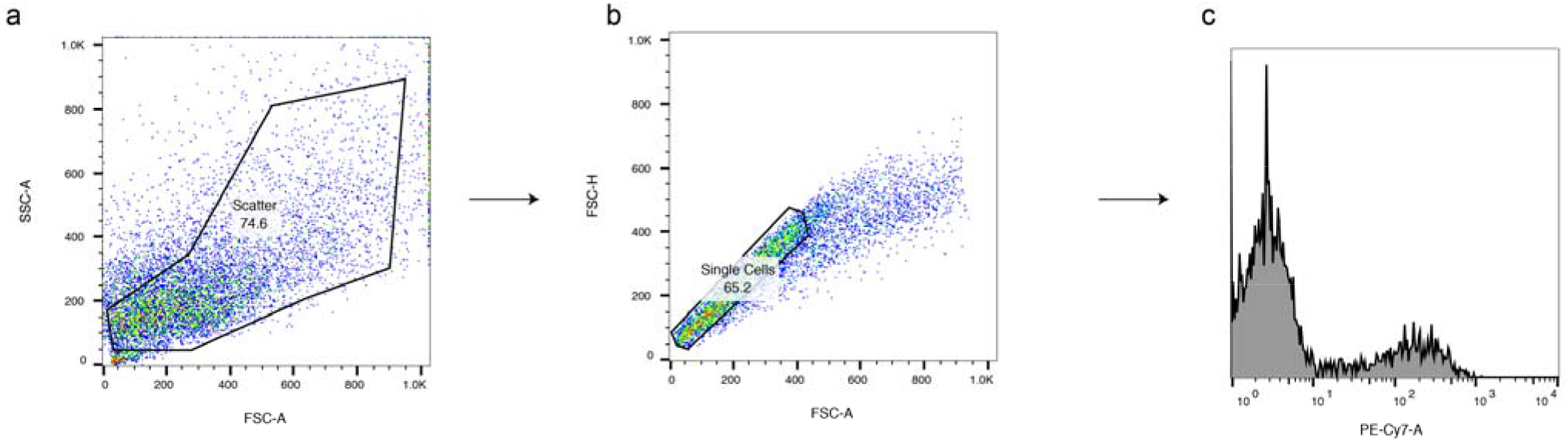
Flow cytometry gating strategy, associated with Fig. 1.

## References

1. Ziegler, C. G. K. et al. SARS-CoV-2 Receptor ACE2 Is an Interferon-Stimulated Gene in Human Airway Epithelial Cells and Is Detected in Specific Cell Subsets across Tissues. Cell 181, 1016–1035.e19 (2020).

2. Chu, H. et al. Comparative Replication and Immune Activation Profiles of SARS-CoV-2 and SARS-CoV in Human Lungs: An Ex Vivo Study With Implications for the Pathogenesis of COVID-19. Clin. Infect. Dis. 71, 1400–1409 (2020).

3. Gutiérrez-Chamorro, L. et al. SARS-CoV-2 infection suppresses ACE2 function and antiviral immune response in the upper respiratory tract of infected patients. bioRxiv 2020.11.18.388850 (2020). doi:10.1101/2020.11.18.388850

4. Blanco-Melo, D. et al. Imbalanced Host Response to SARS-CoV-2 Drives Development of COVID-19. Cell 181, 1036–1045.e9 (2020).

5. Bastard, P. et al. Autoantibodies against type I IFNs in patients with life-threatening COVID-19. Science (80-. ). 370, eabd4585 (2020).

6. Hadjadj, J. et al. Impaired type I interferon activity and inflammatory responses in severe COVID-19 patients. Science (80-. ). 369, 718–724 (2020).

7. Zhang, Q. et al. Inborn errors of type I IFN immunity in patients with life-threatening COVID-19. Science (80-. ). 370, eabd4570 (2020).

8. Samuel, R. M. et al. Androgen Signaling Regulates SARS-CoV-2 Receptor Levels and Is Associated with Severe COVID-19 Symptoms in Men. Cell Stem Cell 27, 876–889.e12 (2020).

9. Daniloski, Z. et al. Identification of Required Host Factors for SARS-CoV-2 Infection in Human Cells. Cell (2020). doi:10.1016/j.cell.2020.10.030

10. Wang, R. et al. Genetic Screens Identify Host Factors for SARS-CoV-2 and Common Cold Coronaviruses. Cell 1–14 (2020). doi:10.1016/j.cell.2020.12.004

11. Schneider, W. M. et al. Genome-Scale Identification of SARS-CoV-2 and Pan-coronavirus Host Factor Networks. Cell 184, 120–132.e14 (2021).

12. Wei, J. et al. Genome-wide CRISPR Screens Reveal Host Factors Critical for SARS-CoV-2 Infection. Cell (2020). doi:10.1016/j.cell.2020.10.028

13. Shi, J. & Vakoc, C. R. The Mechanisms behind the Therapeutic Activity of BET Bromodomain Inhibition. Mol. Cell 54, 728–736 (2014).

14. Fujisawa, T. & Filippakopoulos, P. Functions of bromodomain-containing proteins and their roles in homeostasis and cancer. Nat. Rev. Mol. Cell Biol. 18, 246–262 (2017).

15. Lui, I. et al. Trimeric SARS-CoV-2 Spike interacts with dimeric ACE2 with limited intra-Spike avidity. bioRxiv 2020.05.21.109157 (2020). doi:10.1101/2020.05.21.109157

16. Lan, J. et al. Structure of the SARS-CoV-2 spike receptor-binding domain bound to the ACE2 receptor. Nature 581, 215–220 (2020).

17. Chua, R. L. et al. COVID-19 severity correlates with airway epithelium–immune cell interactions identified by single-cell analysis. Nat. Biotechnol. 38, 970–979 (2020).

18. Tseng, C.-T. K. et al. Apical Entry and Release of Severe Acute Respiratory Syndrome-Associated Coronavirus in Polarized Calu-3 Lung Epithelial Cells. J. Virol. 79, 9470–9479 (2005).

19. Kuchi, S., Gu, Q., Palmarini, M., Wilson, S. J. & Robertson, D. L. Meta-analysis of virus-induced host gene expression reveals unique signatures of immune dysregulation induced by SARS-CoV-2. bioRxiv 2020.12.29.424739 (2020). doi:10.1101/2020.12.29.424739

20. Gilbert, L. a et al. CRISPR-Mediated Modular RNA-Guided Regulation of Transcription in Eukaryotes. Cell 154, 442–51 (2013).

21. Gilbert, L. A. et al. Genome-Scale CRISPR-Mediated Control of Gene Repression and Activation. Cell 159, 647–661 (2014).

22. Horlbeck, M. A. et al. Compact and highly active next-generation libraries for CRISPR-mediated gene repression and activation. Elife 5, 1–20 (2016).

23. Deffieu, M. S. et al. CRISPR-based bioengineering of the Transferrin Receptor revealed a role for Rab7 in the biosynthetic secretory pathway. bioRxiv 2020.01.05.893206 (2020). doi:10.1101/2020.01.05.893206

24. Doroshow, D. B., Eder, J. P. & LoRusso, P. M. BET inhibitors: a novel epigenetic approach. Ann. Oncol. 28, 1776–1787 (2017).

25. Xu, Y. & Vakoc, C. R. Targeting Cancer Cells with BET Bromodomain Inhibitors. Cold Spring Harb. Perspect. Med. 7, a026674 (2017).

26. Filippakopoulos, P. et al. Selective inhibition of BET bromodomains. Nature 468, 1067–1073 (2010).

27. Faivre, E. J. et al. Selective inhibition of the BD2 bromodomain of BET proteins in prostate cancer. Nature 578, 306–310 (2020).

28. Winter, G. E. et al. BET Bromodomain Proteins Function as Master Transcription Elongation Factors Independent of CDK9 Recruitment. Mol. Cell 67, 5–18.e19 (2017).

29. Shi, C. et al. PROTAC induced-BET protein degradation exhibits potent anti-osteosarcoma activity by triggering apoptosis. Cell Death Dis. 10, 815 (2019).

30. Pérez-Bermejo, J. A. et al. SARS-CoV-2 infection of human iPSC-derived cardiac cells predicts novel cytopathic features in hearts of COVID-19 patients. bioRxiv 2020.08.25.265561 (2020). doi:10.1101/2020.08.25.265561

31. Mulay, A. et al. SARS-CoV-2 infection of primary human lung epithelium for COVID-19 modeling and drug discovery. bioRxiv 2020.06.29.174623 (2020). doi:10.1101/2020.06.29.174623

32. Gordon, D. E. et al. A SARS-CoV-2 protein interaction map reveals targets for drug repurposing. Nature 583, 459–468 (2020).

33. Skene, P. J. & Henikoff, S. An efficient targeted nuclease strategy for high-resolution mapping of DNA binding sites. Elife 6, 1–35 (2017).

34. Handoko, L. et al. JQ1 affects BRD2-dependent and independent transcription regulation without disrupting H4-hyperacetylated chromatin states. Epigenetics 13, 410–431 (2018).

35. Wang, S. et al. Target analysis by integration of transcriptome and ChIP-seq data with BETA. Nat. Protoc. 8, 2502–2515 (2013).

36. Au-Yeung, N. & Horvath, C. M. Histone H2A.Z Suppression of Interferon-Stimulated Transcription and Antiviral Immunity Is Modulated by GCN5 and BRD2. iScience 6, 68–82 (2018).

37. Robinot, R. et al. SARS-CoV-2 infection induces the dedifferentiation of multiciliated cells and impairs mucociliary clearance. Nat. Commun. 12, 1–16 (2021).

38. Osterrieder, N. et al. Age-Dependent Progression of SARS-CoV-2 Infection in Syrian Hamsters. Viruses 12, 94301 (2020).

39. Imai, M. et al. Syrian hamsters as a small animal model for SARS-CoV-2 infection and countermeasure development. Proc. Natl. Acad. Sci. U. S. A. 117, 16587–16595 (2020).

40. Sia, S. F. et al. Pathogenesis and transmission of SARS-CoV-2 in golden hamsters. Nature 583, 834–838 (2020).

41. Rosenke, K. et al. Defining the Syrian hamster as a highly susceptible preclinical model for SARS-CoV-2 infection. Emerg. Microbes Infect. 9, 2673–2684 (2020).

42. Gordon, D. E. et al. A SARS-CoV-2 protein interaction map reveals targets for drug repurposing. Nature 583, 459–468 (2020).

43. Ribero, M. S., Jouvenet, N., Dreux, M. & Nisole, S. Interplay between SARS-CoV-2 and the type I interferon response. PLoS Pathog. 16, 1–22 (2020).

44. Lei, X. et al. Activation and evasion of type I interferon responses by SARS-CoV-2. Nat. Commun. 11, 3810 (2020).

45. Xia, H. et al. Evasion of Type I Interferon by SARS-CoV-2. Cell Rep. 33, 108234 (2020).

46. Tian, R. et al. CRISPR Interference-Based Platform for Multimodal Genetic Screens in Human iPSC-Derived Neurons. Neuron 104, 239–255.e12 (2019).

47. Stoner, R., Maures, T. & Conant, D. Methods and Systems for guide RNA Design and Use. (2019).

48. Hsiau, T. et al. Inference of CRISPR Edits from Sanger Trace Data. bioRxiv 251082 (2019). doi:10.1101/251082

49. Adamson, B. et al. A Multiplexed Single-Cell CRISPR Screening Platform Enables Systematic Dissection of the Unfolded Protein Response. Cell 167, 1867–1882.e21 (2016).

50. Glasgow, A. et al. Engineered ACE2 receptor traps potently neutralize SARS-CoV-2. Proc. Natl. Acad. Sci. 117, 28046–28055 (2020).

51. Bolger, A. M., Lohse, M. & Usadel, B. Trimmomatic: A flexible trimmer for Illumina sequence data. Bioinformatics 30, 2114–2120 (2014).

52. Patro, R., Duggal, G., Love, M. I., Irizarry, R. A. & Kingsford, C. Salmon provides fast and bias-aware quantification of transcript expression. Nat. Methods 14, 417–419 (2017).

53. Soneson, C., Love, M. I. & Robinson, M. D. Differential analyses for RNA-seq: transcript-level estimates improve gene-level inferences. F1000Research 4, 1521 (2016).

54. Robinson, M. D., McCarthy, D. J. & Smyth, G. K. edgeR: a Bioconductor package for differential expression analysis of digital gene expression data. Bioinformatics 26, 139–140 (2010).

55. Eisen, M. B., Spellman, P. T., Brown, P. O. & Botstein, D. Cluster analysis and display of genome-wide expression patterns. Proc. Natl. Acad. Sci. 95, 14863–14868 (1998).

56. Saldanha, A. J. Java Treeview--extensible visualization of microarray data. Bioinformatics 20, 3246–3248 (2004).

57. Skene, P. J., Henikoff, J. G. & Henikoff, S. Targeted in situ genome-wide profiling with high efficiency for low cell numbers. Nat. Protoc. 13, 1006–1019 (2018).

58. Meers, M. P., Tenenbaum, D. & Henikoff, S. Peak calling by Sparse Enrichment Analysis for CUT&RUN chromatin profiling. Epigenetics Chromatin 12, 42 (2019).

